# Genes Affecting Vocal and Facial Anatomy Went Through Extensive Regulatory Divergence in Modern Humans

**DOI:** 10.1101/106955

**Authors:** David Gokhman, Malka Nissim-Rafinia, Lily Agranat-Tamir, Genevieve Housman, Raquel García-Pérez, Esther Lizano, Olivia Cheronet, Swapan Mallick, Maria A. Nieves-Colón, Heng Li, Songül Alpaslan-Roodenberg, Mario Novak, Hongcang Gu, Jason M. Osinski, Manuel Ferrando-Bernal, Pere Gelabert, Iddi Lipende, Deus Mjungu, Ivanela Kondova, Ronald Bontrop, Ottmar Kullmer, Gerhard Weber, Tal Shahar, Mona Dvir-Ginzberg, Marina Faerman, Ellen E. Quillen, Alexander Meissner, Yonatan Lahav, Leonid Kandel, Meir Liebergall, María E. Prada, Julio M. Vidal, Richard M. Gronostajski, Anne C. Stone, Benjamin Yakir, Carles Lalueza-Fox, Ron Pinhasi, David Reich, Tomas Marques-Bonet, Eran Meshorer, Liran Carmel

## Abstract

Regulatory changes are broadly accepted as key drivers of phenotypic divergence. However, identifying regulatory changes that underlie human-specific traits has proven very challenging. Here, we use 63 DNA methylation maps of ancient and present-day humans, as well as of six chimpanzees, to detect differentially methylated regions that emerged in modern humans after the split from Neanderthals and Denisovans. We show that genes affecting the face and vocal tract went through particularly extensive methylation changes. Specifically, we identify widespread hypermethylation in a network of face- and voice-affecting genes (*SOX9*, *ACAN*, *COL2A1*, *NFIX* and *XYLT1*). We propose that these repression patterns appeared after the split from Neanderthals and Denisovans, and that they might have played a key role in shaping the modern human face and vocal tract.

## Introduction

The advent of high-coverage ancient genomes of archaic humans (Neanderthal and Denisovan) introduced the possibility to identify the genetic basis of some unique modern human traits^1^. A common approach is to carry out sequence comparisons and detect non-neutral sequence changes. However, out of ∼30,000 substitutions and indels that reached fixation in modern humans, less than 100 directly alter amino acid sequence^1^, and as of today, our ability to estimate the biological effects of the remaining ∼30,000 noncoding changes is very restricted. Whereas many of them are probably nearly neutral, many others may affect gene function, especially those in regulatory regions such as promoters and enhancers. Such regulatory changes may have a sizeable impact on human evolution, as alterations in gene regulation are thought to underlie much of the phenotypic variation between closely related groups^2^. Because of the limited ability to interpret noncoding variants, direct examination of regulatory layers such as DNA methylation has the potential to enhance our understanding of the evolutionary origin of human-specific traits far beyond what can be achieved using sequence comparison alone^3^.

In order to gain insight into the regulatory changes that underlie human evolution, we previously developed a method to reconstruct DNA methylation maps of ancient genomes based on analysis of patterns of damage to ancient DNA^4^. We used this method to reconstruct the methylomes of a Neanderthal and a Denisovan, which were then compared to a partial methylation map of a present-day osteoblast cell line. However, the ability to identify differentially methylated regions (DMRs) between the human groups was constrained by the incomplete reference map (providing methylation information for ∼10% of CpG sites), differences in outputs of sequencing platforms, lack of an outgroup, and a restricted set of skeletal samples (see Methods).

To study the evolutionary dynamics of DNA methylation along the hominin tree on a larger scale, we establish here the most comprehensive assembly to date of skeletal DNA methylation maps from modern humans, archaic humans, and chimpanzees. Using these maps, we identify 588 genes whose methylation state is unique to modern human. We then analyze the function of these genes by investigating their known anatomical effects, and validate this using over 50 orthogonal tests and controls. We find that the most extensive DNA methylation changes are observed in genes that affect vocal and facial anatomy, and that this trend is unique to modern humans.

## Results

We reconstructed ancient DNA methylation maps of eight individuals: in addition to the previously published Denisovan and Altai Neanderthal methylation maps^4^, we reconstructed the methylomes of the Vindija Neanderthal (∼52 thousand years ago, kya)^5^, and three anatomically modern humans: the Ust’-Ishim individual (∼45 kya, Western Siberia)^6^, the Loschbour individual (∼8 kya, Luxemburg)^7^, and the Stuttgart individual (∼7 kya, Germany)^7^. We also sequenced to high-coverage and reconstructed the methylomes of the La Braña 1 individual from Spain (∼8 kya, 22x) (which was previously sequenced to low-coverage^8^) and an individual from Barçın Höyük, Western Anatolia, Turkey (I1583, ∼8.5 kya, 24x), which was previously sequenced using a capture array^9^.

To this set we added 52 publicly available partial bone methylation maps from present-day individuals, produced using 450K methylation arrays (see Methods). To obtain full present-day bone maps, we produced whole-genome bisulfite sequencing (WGBS) methylomes from the femur bones of two individuals (Bone1 and Bone2). Hereinafter, ancient and present-day modern humans are collectively referred to as *anatomically modern humans* (AMHs), while the Neanderthal and Denisovan are referred to as *archaic humans*. As an outgroup, we produced methylomes of six chimpanzees: one WGBS, one reduced representation bisulfite sequencing (RRBS) and four 850K methylation arrays. Together, these data establish a unique and comprehensive platform to study DNA methylation dynamics in recent human evolution (Extended Data Table 1).

## Identification of DMRs

We developed a DMR-detection method for ancient methylomes, which accounts for potential noise introduced during reconstruction, as well as differences in coverage and deamination rates (Extended Data Fig. 1). To minimize the number of false positives and to identify DMRs that are most likely to have a regulatory effect, we applied a strict threshold of >50% difference in methylation across a minimum of 50 CpGs. This also filters out environmentally-induced DMRs which typically show small methylation differences and limited spatial scope^10^. Using this method, we identified 9,679 regions overall that showed methylation differences between any of the high-quality representative methylomes of the Denisovan, the Altai Neanderthal, and the Ust’-Ishim anatomically modern human. These regions do not necessarily represent evolutionary differences between the human groups. Rather, many of them could be attributed to factors separating the three individuals (e.g., Ust’-Ishim is a male whereas the archaic humans are females), or to variability within populations. To minimize such effects, we used the 59 additional human maps to filter out regions where variability in methylation is detected. We adopted a conservative approach, whereby we take only loci where methylation in one hominin group is found completely outside the range of methylation in the other groups (Fig. 1a).

**Figure 1.**
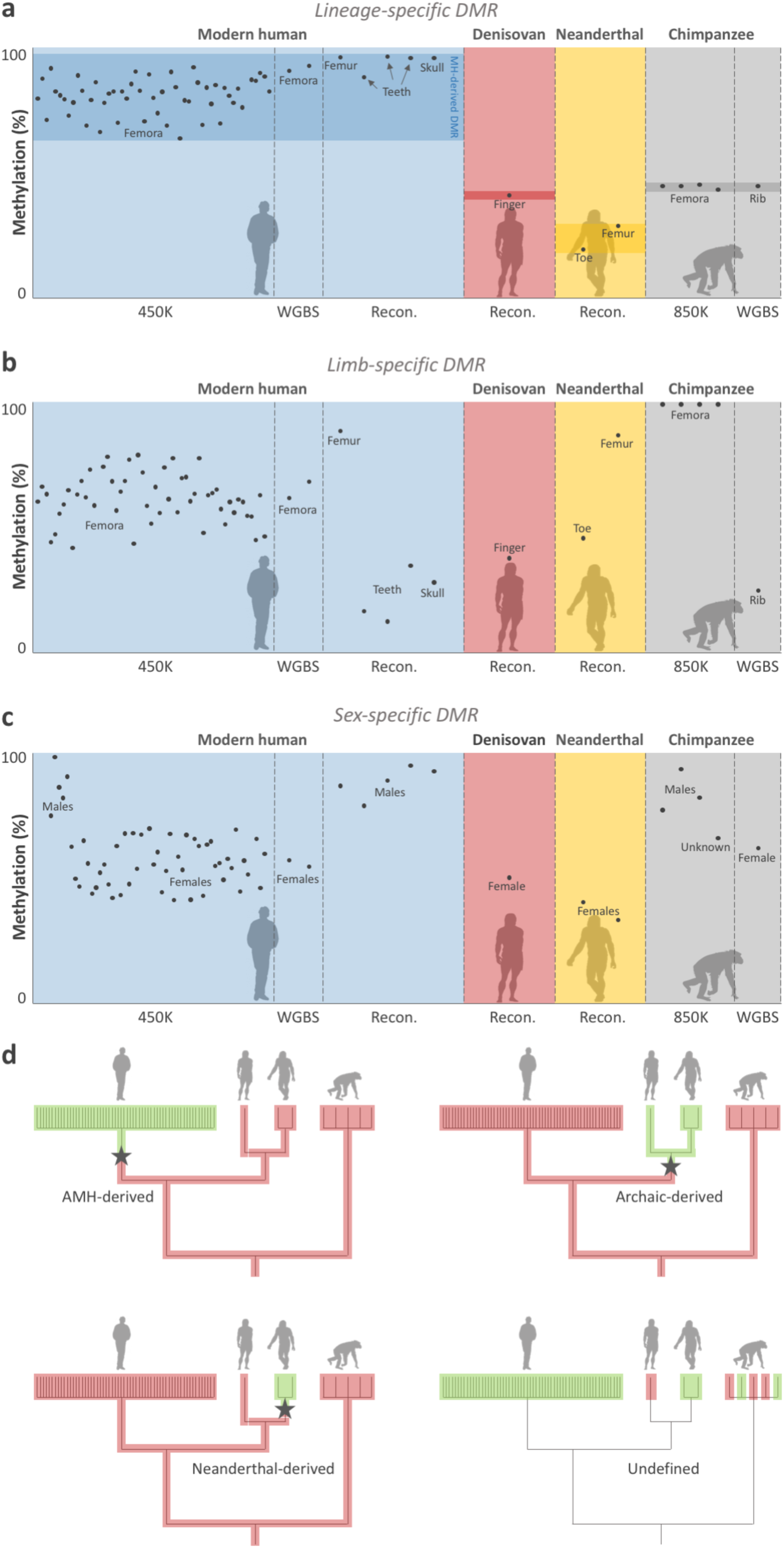
Variability filtering and lineage assignment. **a.** Methylation levels across AMH, Denisovan, Neanderthal, and chimpanzee samples in DMR#278 (chr4:38,014,896-38,016,197). This is an example of a lineage-specific DMR, defined as a locus in which all samples of a group are found outside the range of methylation in the other groups. Chimpanzee samples were used during the following step of lineage assignment. **b.** A putative limb-specific DMR (chr3:14,339,371-14,339,823) which was removed from the analysis, as it does not comply with our definition of lineage-specific DMRs. Femur, toe, and finger samples are hypermethylated compared to other skeletal elements. Toe and finger are found at the bottom range of limb samples, suggesting some variation in this locus within limb samples too. **c.** A putative sex-specific DMR (chr3:72,394,336-72,396,901) which was removed from the analysis. Males are hypermethylated compared to females. **d.** Lineage assignment using chimpanzee samples. Each bar at the tree leaves represents a sample. Methylation levels are marked with red and green, representing methylated and unmethylated samples, respectively. Only DMRs that passed the previous variability filtering steps were analyzed. The lineage where the methylation change has likely occurred (by parsimony) is marked by a star. Branch lengths are not scaled.

Importantly, our samples come from both sexes, from individuals of various ages and ancestries, from sick and healthy individuals, and from a variety of skeletal parts (femur, skull, phalanx, tooth, and rib). Hence, this procedure is expected to account for these potentially confounding factors, and the remaining DMRs are expected to represent true evolutionary differences (Fig. 1a-c, Extended Data Fig. 1, see Methods). This step resulted in a set of 7,649 DMRs that discriminate between the human groups, which we ranked according to their significance level. Next, using the chimpanzee samples, we were able to determine for 2,825 of these DMRs the lineage where the methylation change occurred (Fig. 1d). Of these DMRs, 873 are AMH-derived, 939 are archaic-derived, 443 are Denisovan-derived, and 570 are Neanderthal-derived (Fig. 2a, Extended Data Fig. 1, Extended Data Table 2). To study the derived biology of AAMHs, and to focus on DMRs that are based on the most extensive set of maps, we concentrated on the 873 AMH-derived DMRs. We found that these DMRs are located 58x closer to AMH-derived sequence changes than expected by chance (0.092 Mb vs. a median of 5.3 Mb, *P* < 10^-5^, permutation test, Fig. 2b). This suggests that some of the methylation changes might have been driven by cis-regulatory sequence variants that arose along the AMH lineage.

**Figure 2.**
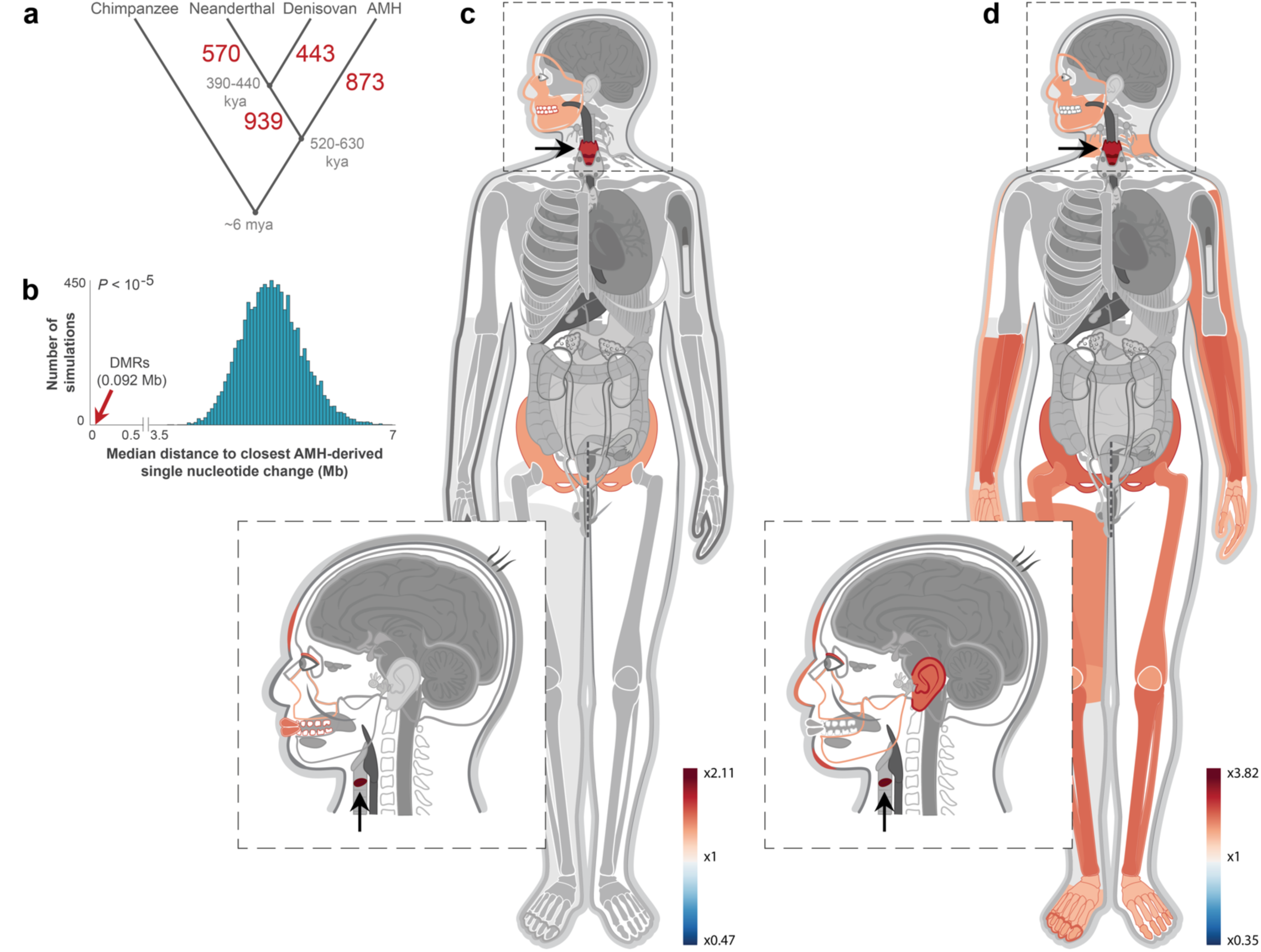
Genes affecting voice and face are the most over-represented within AMH-derived DMRs. **a.** The number of DMRs that emerged along each of the human branches. Divergence times are in thousands of years ago (kya). **b**. Distribution of median distances (turquoise) of DMRs to randomized single nucleotide changes that separate AMHs from archaic humans and chimpanzees. Genomic positions of single nucleotide changes were allocated at random. This was repeated 10,000 times. Red arrow marks the observed distance of DMRs, showing that they tend to be significantly closer to AMH-derived single nucleotide changes than expected by chance. This suggests that some of these sequence changes might have driven the changes in methylation. **c.** A heat map representing the level of enrichment of each anatomical part within the AMH-derived DMRs. Only body parts that are significantly enriched (FDR < 0.05) are colored. Three skeletal parts are significantly over-represented: the face, pelvis, and larynx (voice box, marked with arrows). **d.** Enrichment levels of anatomical parts within the most significant (top quartile, *Q* statistic) AMH-derived DMRs, showing a more pronounced enrichment of genes affecting vocal and facial anatomy.

### Face and voice-affecting genes are derived in AMHs

We defined differentially methylated genes (DMGs) as genes that overlap at least one DMR along their body or in their promoter, up to 5 kb upstream. The 873 AMH-derived DMRs are linked to 588 AMH-derived DMGs (Extended Data Table 2). To gain insight into the function of these DMGs, we first analyzed their gene ontology (GO). As expected from genes that show differential methylation in the skeleton between human groups, AMH-derived DMGs are enriched with terms associated with the skeleton (e.g., *endochondral bone morphogenesis, trabecula morphogenesis,* and *palate development*). Also notable are terms associated with the skeletal muscular, cardiovascular, and nervous systems (Extended Data Table 3).

To acquire a more precise understanding of the possible functional consequences of these DMGs, we used Gene ORGANizer, which links human genes to the organs they phenotypically affect^11^. Unlike tools that use GO terms or RNA expression data, Gene ORGANizer is based entirely on curated gene-disease and gene-phenotype associations from monogenic diseases. It relies on direct phenotypic observations in human patients whose conditions result from known gene perturbations. Using Gene ORGANizer, we found 11 organs that are over-represented within the 588 AMH-derived DMGs, eight of which are skeletal parts that can be divided into three regions: the face, larynx (voice box), and pelvis (Fig. 2c, Extended Data Table 4). The strongest enrichment was observed in the laryngeal region (x2.11 and x1.68, FDR = 0.017 and 0.048, for the vocal folds (vocal cords) and larynx, respectively), followed by facial and pelvic structures, including the teeth, forehead, jaws, and pelvis. Interestingly, the face and pelvis are considered the most morphologically divergent regions between Neanderthals and AMHs^12^ and our results reflect this divergence through gene regulation changes. To gain orthogonal evidence for the enrichment of the larynx and face within these AMH-derived DMGs, we carried out a number of additional analyses: First, we analyzed gene expression patterns and found that the supralaryngeal vocal tract (the pharyngeal, oral, and nasal cavities, where sound is filtered to specific frequencies) is the most enriched body part (1.7x and 1.6x, FDR = 5.6 x 10^-6^ and FDR = 7.3 x 10^-7^, for the pharynx and larynx, respectively, Extended Data Table 3). Second, 44 of the AMH-derived DMRs overlap previously reported putative enhancers of human craniofacial developmental genes (5.1x compared to expected, *P* < 10^-4^, permutation test)^13, 14^. Third, *Palate development* is the third most enriched GO term among AMH-derived DMGs (Extended Data Table 3). Fourth, DMGs significantly overlap genes associated with craniofacial features in the GWAS catalog^15^ (*P* = 3.4 x 10^-4^, hypergeometric test).

To test whether this enrichment remains if we take only the most confident DMRs, we limited the analysis to DMGs where the most significant DMRs are found (top quartile, *Q* statistic). Here, the over-representation of voice-affecting genes is even more pronounced (2.82x and 2.26x, for vocal folds and larynx, respectively, FDR = 0.028 for both, Fig. 2d, Extended Data Table 4). Hereinafter, we refer to genes as affecting an organ if they have been shown to have a phenotypic effect on that organ in some or all patients where this gene is dysfunctional. Next, we reasoned that skeleton-associated genes might be over-represented in analyses that compare bone DNA methylation maps, hence introducing potential biases. To test whether this enrichment might explain the over-representation of the larynx, face, and pelvis, we compared the fraction of genes affecting these organs within all skeletal genes to their fraction within the skeletal genes in the AMH-derived DMGs. We found that genes affecting the face, larynx, and pelvis are significantly over-represented even within skeletal AMH-derived DMGs (*P* = 1.0 x 10^-5^, *P* = 1.3 x 10^-3^, *P* = 2.1 x 10^-3^, *P* = 0.03, for vocal folds, larynx, face, and pelvis, respectively, hypergeometric test). Additionally, using a permutation test, we found that the enrichment levels within AMH-derived DMGs are significantly higher than expected by chance for the laryngeal and facial regions, but not for the pelvis (*P* = 8.0 x 10^-5^, *P* = 3.6 x 10^-3^, *P* = 8.2 x 10^-4^, and *P* = 0.115, for vocal folds, larynx, face and pelvis, respectively, Extended Data Fig. 2b-e, see Methods). Thus, we found that the enrichment in the facial and laryngeal regions is not a by-product of a general enrichment in skeletal parts, and we hereinafter focus on genes affecting these two regions.

Finally, we ruled out the options that our DMR-detection algorithm, number of samples, filtering process or biological factors such as gene length, cellular composition, pleiotropy or developmental stage might underlie the enrichment of these organs (see Methods).

Perhaps most importantly, none of the other branches shows enrichment of the larynx or the vocal folds; Neanderthal- and Denisovan-derived DMGs show no significant enrichment in any organ, and archaic-derived DMGs are over-represented in the jaws, lips, limbs, scapulae, and spinal column, but not in the larynx or vocal folds (Extended Data Fig. 2f, Extended Data Table 4). In addition, DMRs that separate chimpanzees from all humans (archaic and modern, Extended Data Table 2) do not show enrichment of genes affecting the larynx or face, compatible with the notion that this trend emerged along the AMH lineage.

Taken together, we conclude that DMGs that emerged along the AMH lineage are uniquely enriched in genes affecting the voice and face, and that this is unlikely to be an artifact of (a) inter-individual variability resulting from age, sex, disease, or bone type; (b) significance level of DMRs; (c) the reconstruction or DMR-detection processes; (d) number of samples used; (e) pleiotropic effects; (f) the types of methylation maps used; (g) the comparison of skeletal methylomes; (h) gene length distribution; or (i) biological factors such as cellular composition and developmental state.

Our analyses identified 56 DMRs in genes affecting the facial skeleton, and 32 in genes affecting the laryngeal skeleton. The face-affecting genes are known to shape mainly the protrusion of the lower and midface, the size of the nose, and the slope of the forehead. Interestingly, these traits are considered some of the most derived between Neanderthals and AMHs^12^. The larynx-affecting genes have been shown to underlie various phenotypes in patients, ranging from slight changes to the pitch and hoarseness of the voice, to a complete loss of speech ability^11^ (Extended Data Table 5). These phenotypes were shown to be driven primarily by alterations to the laryngeal and vocal tract skeleton. Methylation patterns in differentiated cells are often established during earlier stages of development, and the closer two tissues are developmentally, the higher the similarity between their methylation maps^3, 16, 17^. This is also evident in the fact that DMRs identified between species in one tissue often exist in other tissues as well^16^. Importantly, the laryngeal skeleton, and particularly the arytenoid cartilage to which the vocal folds are anchored, share an origin from the somatic layer of the lateral plate mesoderm with the cartilaginous tissue of the limb bones prior to their ossification. Thus, it is likely that many of the DMRs identified here between limb samples also exist in their closest tissue – the laryngeal skeleton. This is further supported by the observation that these DMGs are consistent across all examined skeletal samples, including skull, femur, rib, tibia, and tooth. Furthermore, we directly measured methylation levels in a subset of the DMRs in primary chondrocytes and show that their patterns extend to these cells as well (see below).

### Extensive methylation changes within face and voice-affecting genes

The results above suggest that methylation levels in many face- and voice-affecting genes have changed in AMHs since the split from archaic humans, but they do not provide information on the extent of changes within each gene. To do so, we scanned the genome in windows of 100 kb and computed the fraction of CpGs which are differentially methylated in AMHs (hereinafter, AMH-derived CpGs). We found that the extent of changes within voice-affecting DMGs is most profound, more than 2x compared to other DMGs (0.132 vs. 0.055, FDR = 2.3 x 10^-3^, *t*-test, Extended Data Table 6). Face-affecting DMGs also present high density of AMH-derived CpGs (0.079 vs. 0.055, FDR = 2.8 x 10^-3^). In archaic-derived DMGs, on the other hand, the extent of changes within voice- and face-affecting genes is not different than expected (FDR = 0.99, Extended Data Table 6). To control for possible biases, we repeated the analysis using only the subset of DMRs in genes affecting the skeleton. Here too, we found that voice-affecting AMH-derived DMGs present the highest density of changes (2.5x for vocal folds, 2.4x for larynx, FDR = 1.4 x 10^-3^ for both, Extended Data Table 6), and face-affecting DMGs also exhibit a significantly elevated density of changes (1.4x, FDR = 0.04).

We also found that compared to other AMH-derived DMRs, DMRs in voice- and face-affecting genes tend to be 40% closer to candidate positively selected loci in AMHs^18^ (*P* < 10^-4^, permutation test).

Strikingly, when ranking DMGs according to the fraction of AMH-derived CpGs, all top five skeleton-related DMGs (*ACAN*, *SOX9*, *COL2A1*, *XYLT1*, and *NFIX*) are known to affect lower and midfacial protrusion, as well as the voice^11, 19^ (Fig. 3a,b, Extended Data Fig. 2g). This is particularly surprising considering that genome-wide, less than 2% of genes (345) are known to affect the voice, ∼3% of genes (726) are known to affect lower and midfacial protrusion, and less than 1% (182) are known to affect both^11, 19^.

**Figure 3.**
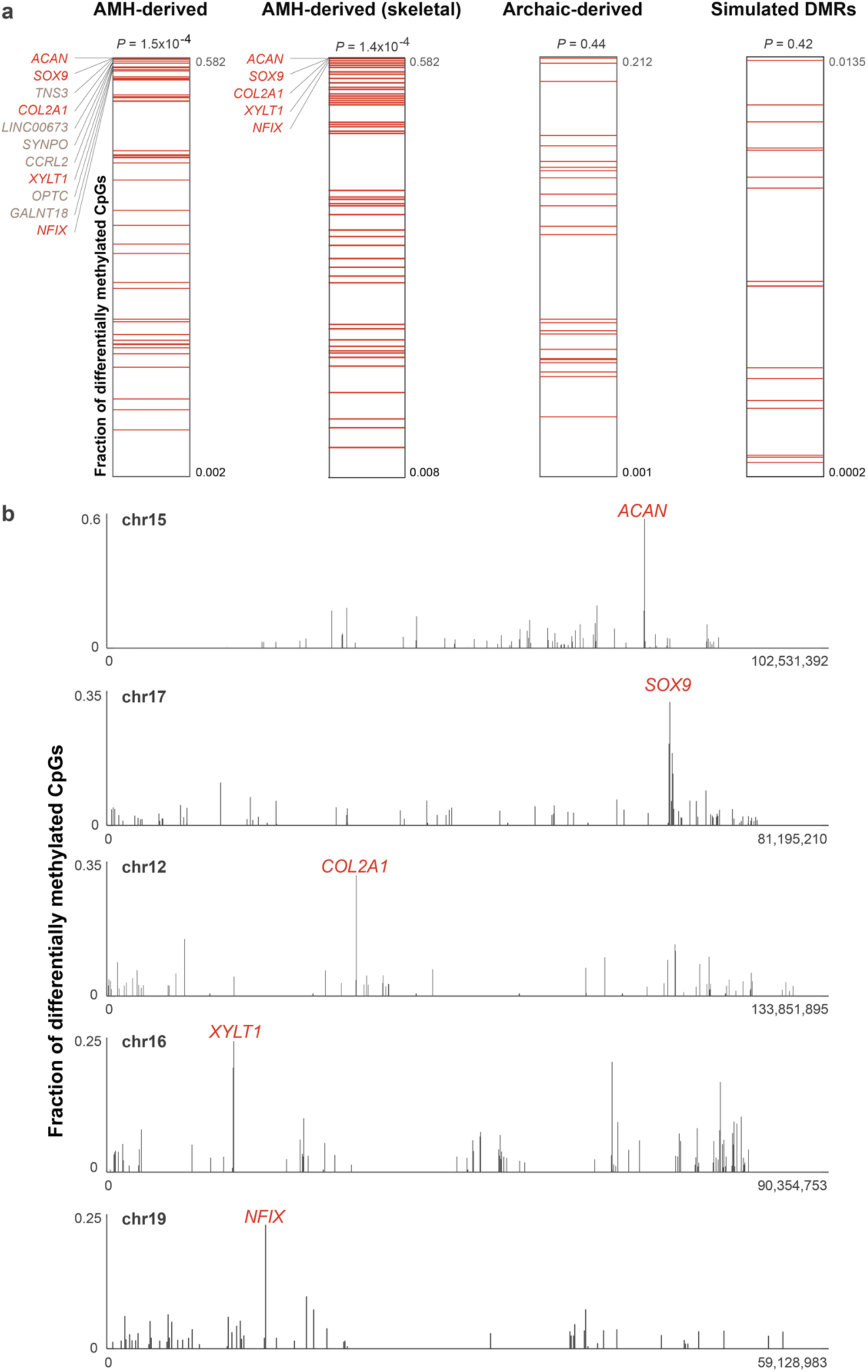
The extent of differential methylation is highest among genes affecting the voice. **a.** Within each lineage, the fraction of differentially methylated CpGs was computed as the number of derived CpGs per 100 kb centered around the middle of each DMR. Genes were ranked according to the fraction of derived CpG positions within them. Genes affecting the voice are marked with red lines. AMH-derived DMRs in voice-affecting genes tend to be ranked significantly higher. Although these genes comprise less than 2% of the genome, three of the top five AMH-derived DMRs, and all top five skeleton-related AMH-derived DMRs are in genes that affect the voice. In archaic-derived DMRs and in simulated DMRs, voice-affecting genes do not show higher ranking compared to the rest of the DMGs. **b.** The fraction of differentially methylated CpGs along the five chromosomes containing *ACAN*, *SOX9*, *COL2A1*, *XYLT1*, and *NFIX*. In each of these chromosomes, the most extensive changes are found within these genes. All five genes control facial projection and the development of the larynx.

The three skeletal DMGs with the highest density of AMH-derived CpGs are the extra-cellular matrix genes *ACAN* and *COL2A1,* and their key regulator *SOX9*, which together form a network that regulates skeletal growth, the transition from cartilage to bone, and spatio-temporal patterning of skeletal development, including the facial and laryngeal skeleton in humans^19, 20^ and mouse^21^. *SOX9* was also shown to be one of the top genes underlying variation in craniofacial morphology within-AMHs^22^. *SOX9* is regulated by a series of upstream enhancers identified in mouse and human^23^. In human skeletal samples, hypermethylation of the *SOX9* promoter was shown to down-regulate its activity, and consequently its targets^24^. This was also demonstrated repeatedly in non-skeletal tissues of human^25, 26^ and mouse^27, 28^. We found substantial hypermethylation in AMHs in the following regions: (a) the *SOX9* promoter; (b) seven of its proximal and distal skeletal and skeletal progenitor enhancers^23^; (c) the targets of SOX9: *ACAN* (DMR #80) and *COL2A1* (DMR #1, the most significant AMH-derived DMR, which spans 32kb and covers almost the entire *COL2A1* gene, from its 1^st^ intron to its 54^th^ exon and 3’UTR region); and (d) an upstream lincRNA (*LINC02097*). Notably, regions (a), (b), and (d) overlap the longest DMR on the AMH-derived DMR list, spanning 35,910 bp (DMR #11, Fig. 4).

**Figure 4.**
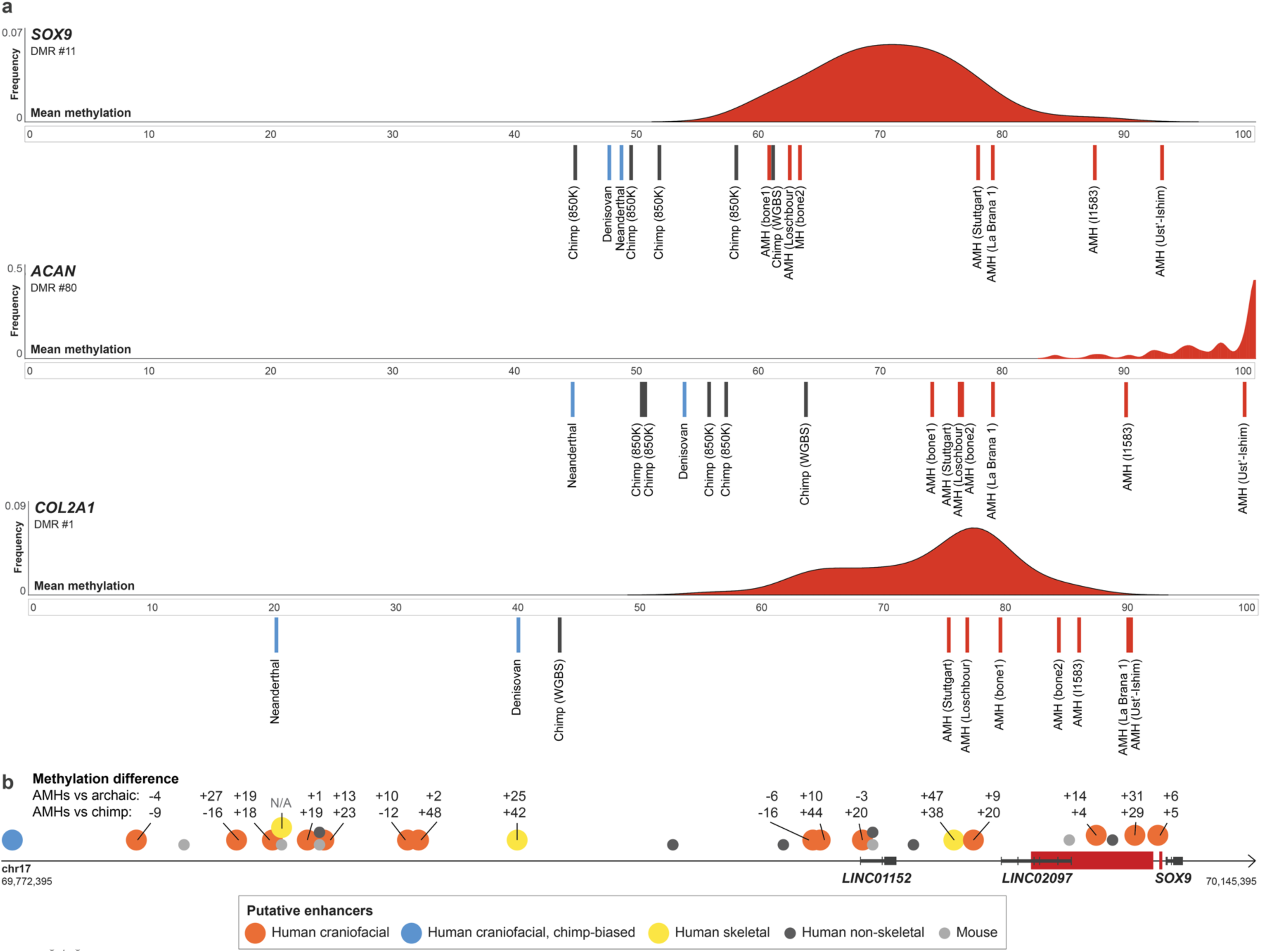
Hypermethylation of *SOX9*, *ACAN*, and *COL2A1* in AMHs. **a.** Methylation levels in the AMH-derived DMRs in *SOX9*, *ACAN*, and *COL2A1*. AMH samples are marked with red lines, archaic human samples are marked with blue lines and chimpanzee samples are marked with grey lines. The distribution of methylation across 52 AMH samples (450K methylation arrays) is presented in red. **b.** *SOX9* and its upstream regulatory elements. AMH-derived DMRs are marked with red rectangles. Previously identified putative enhancers are marked with circles. Numbers above skeletal enhancers show the difference in mean bone methylation between AMHs and archaic humans (top) and between AMHs and chimpanzee (bottom). Across almost all *SOX9* enhancers, AMHs are hypermethylated compared to archaic humans and the chimpanzee.

Additionally, a more distant putative enhancer, located 345 kb upstream of *SOX9*, was shown to bear strong active histone modification marks in chimpanzee craniofacial progenitor cells; whereas, in humans these marks are almost absent (∼10x lower than chimpanzee, suggesting down-regulation, Fig. 4b)^13^. Importantly, human and chimpanzee non-skeletal tissues (i.e., brain and blood) exhibit very similar methylation patterns in these genes, suggesting the DMRs are skeleton-specific. Finally, the amino acid sequence coded by each of these genes is identical across the hominin groups^1^, suggesting that the observed changes are purely regulatory. Together, these observations support the idea that *SOX9* became down-regulated in AMH skeletal tissues, likely followed by down-regulation of its targets: *ACAN* and *COL2A1*. *XYLT1*, the 4^th^ highest skeleton-related DMG, is an enzyme involved in the synthesis of glycosaminoglycan. Loss-of-function mutations, hypermethylation of the gene and its consequent reduced expression underlie the Desbuquois dysplasia skeletal syndrome, which was shown to affect the cartilaginous structure of the larynx, and drive a retraction of the face^29, 30^. Very little is known about *XYLT1* regulation, but interestingly, in zebrafish it was shown to be bound by SOX9 [^31^].

To quantitatively investigate the potential phenotypic consequences of these DMGs, we tested what fraction of their known phenotypes are also known as traits that differ between modern and archaic humans. We found that four of the top five most differentially methylated genes (*XYLT1*, *NFIX*, *ACAN*, and *COL2A1*) are in the top 100 genes with the highest fraction of divergent traits between Neanderthals and AMHs. Remarkably, *COL2A1*, the most divergent gene in its methylation patterns, is also the most divergent in its phenotypes: no other gene in the genome is associated with as many divergent traits between modern humans and Neanderthals (63 traits, Extended Data Table 7, see Methods). This suggests that these extensive methylation changes are possibly linked to phenotypic divergence between archaic and AMHs.

### *NFIX* methylation patterns suggest downregulation in AMHs

In order to investigate how methylation changes affect expression levels, we scanned the DMRs to identify those whose methylation levels are strongly correlated with expression across 22 human tissues^32^. We found 90 such AMH-derived DMRs (FDR < 0.05, Extended Data Table 2). DMRs in voice-affecting genes are significantly more likely to be correlated with expression compared to other DMRs (2.05x, *P* = 6.65 x 10^-4^, hypergeometric test). Particularly noteworthy is *NFIX*, one of the most derived genes in AMHs (ranked 5^th^ among DMGs affecting the skeleton, Fig. 3a,b)*. NFIX* contains two DMRs (#24 and #167, Fig. 5a), and in both, methylation levels are tightly linked with expression (correlation of 81.7% and 73.8%, FDR = 3.5 x 10^-6^ and 8.6 x 10^-5^, respectively, Fig. 5b). In fact, *NFIX* is one of the top ten DMGs with the most significant correlation between methylation and expression in human. The association between *NFIX* methylation and expression was also shown previously across several mouse tissues^33, 34^. To further examine this, we investigated a dataset of DNMT3A-induced methylation of human MCF-7 cells. Forced induction of methylation in this study was sufficient to repress *NFIX* expression by over 50%, placing *NFIX* as one of the genes whose expression is most affected by hypermethylation^35^ (ranked in the 98^th^ percentile, FDR = 1.28 x 10^-6^). We further validated the hypermethylation of *NFIX* across the skeleton by comparing four human cranial samples to four chimpanzee cranial samples through bisulfite-PCR (*P* = 0.01, Extended Data Fig. 3, Extended Data Table 1, see Methods). Together, these findings suggest that the observed hypermethylation of *NFIX* in AMHs reflects down-regulation that emerged along the AMH lineage. Indeed, we found that *NFIX*, as well as *SOX9, ACAN*, *COL2A1*, and *XYLT1* are hypermethylated in human femora compared to baboon^36^ (*P* = 1.4 x 10^-5^ and *P* = 8.1 x 10^-9^, compared to baboon femora bone and cartilage, respectively, *t*-test). Also, all five genes show significantly reduced expression in humans compared to mice (Fig. 5c). Taken together, these observations suggest that DNA methylation is a primary mechanism in the regulation of *NFIX*, and serves as a good proxy for its expression. Interestingly, NFI proteins were shown to bind the upstream enhancers of *SOX9* [^37^], hence suggesting a possible mechanism to the simultaneous changes in the five top genes we report.

**Figure 5.**
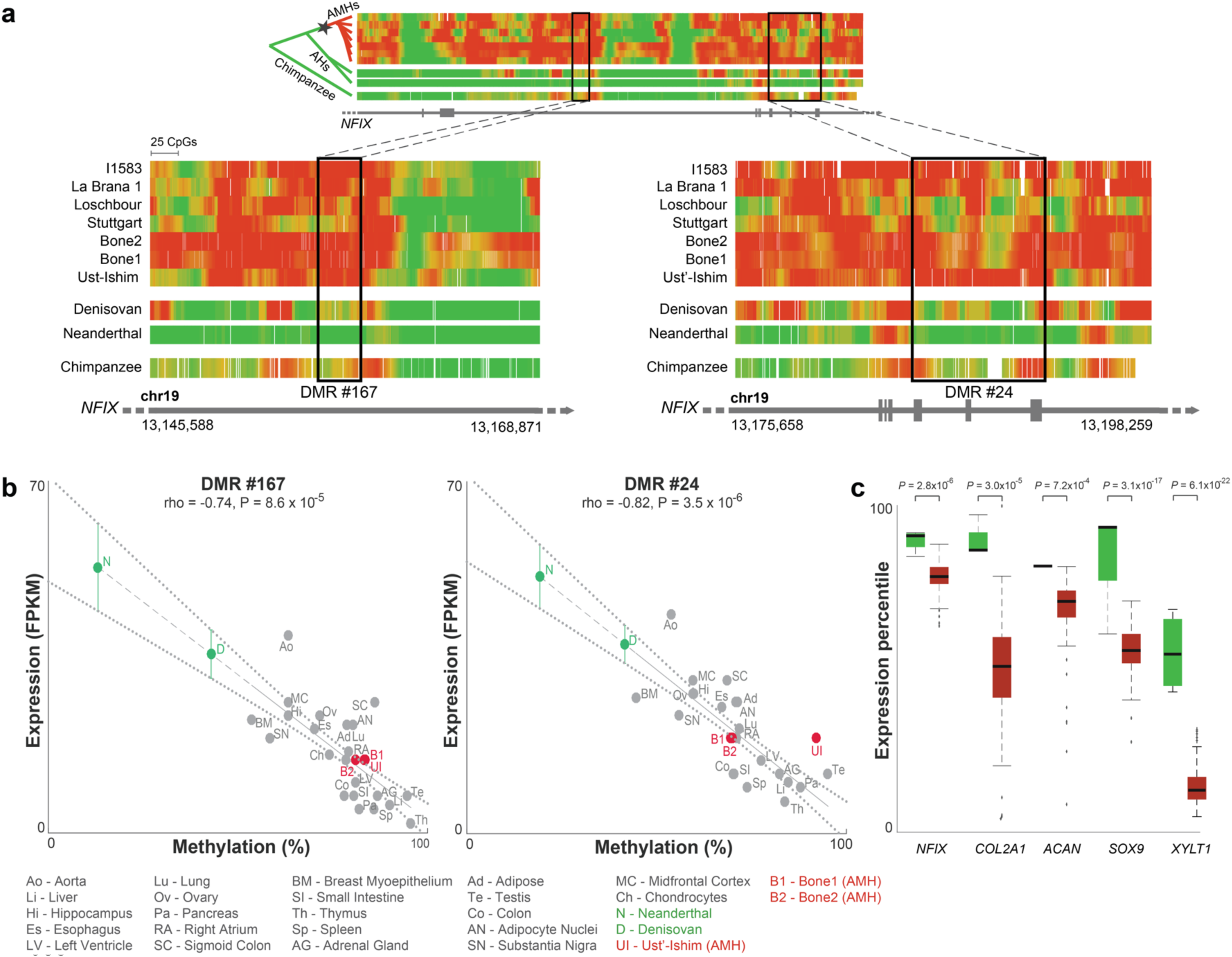
*NFIX* became down-regulated after the split from archaic humans. **a.** Methylation levels along *NFIX*, color-coded from green (unmethylated) to red (methylated). Methylation levels around the two AMH-derived DMRs (#24 and #167) are shown in the zoomed-in panels. These two DMRs represent the regions where the most significant methylation changes are observed, but hypermethylation of *NFIX* in AMHs can be seen throughout the entire gene body. Chimpanzee and present-day samples were smoothed using the same sliding window as in ancient samples to allow easier comparison. The inferred schematic regulatory evolution of *NFIX* is shown using a phylogenetic tree to the left of the top panel. Star marks the shift in methylation from unmethylated (green) to methylated (red). **b**. Methylation levels in DMRs #167 and #24 vs. expression levels of *NFIX* across 22 AMH tissues (grey). In both DMRs, higher methylation is significantly associated with lower expression of *NFIX*. Ust’-Ishim, Bone1 and Bone2 methylation levels (red) are plotted against mean NFIX expression across 13 osteoblast lines. Neanderthal and Denisovan methylation levels (green) are plotted against the predicted expression levels, based on the extrapolated regression line (dashed). Standard errors are marked with dotted lines. The Neanderthal and Denisovan are expected to have higher *NFIX* expression levels. **c.** Expression levels of *NFIX, COL2A1, ACAN, SOX9* and *XYLT1* in AMHs are reduced compared to mice. Box plots present 89 human samples (red) and four mouse samples (green) from appendicular bones (limbs and pelvis). Expression levels were converted to percentiles based on the level of gene expression compared to the rest of the genome in each sample.

## Discussion

We have shown here that genes affecting vocal and facial anatomy went through extensive methylation changes in recent AMH evolution, after the split from Neanderthals and Denisovans. The extensive methylation changes are manifested both in the number of divergent genes and in the extent of changes within each gene. Notably, the DMRs we report capture substantial methylation changes (over 50% between at least one pair of human groups), span thousands or tens of thousands of bases, and cover promoters and enhancers. Many of these methylation changes are tightly linked with changes in expression. We particularly focused on changes in the regulation of the five most derived skeletal genes on the AMH lineage: *SOX9*, *ACAN*, *COL2A1*, *XYLT1*, and *NFIX*, whose downregulation was shown to underlie a retracted face, as well as changes to the structure of the larynx^20, 29, 38–41^. The results we report, which are based on ancient DNA methylation patterns, provide novel means to analyze the genetic mechanisms that underlie the evolution of the human face and vocal tract.

Humans are distinguished from other apes in their unique capability to communicate through speech. This capacity is attributed not only to neural changes, but also to structural alterations to the vocal tract^42^. The relative roles of anatomy vs. cognition in our speech skills are still debated^43^, and some propose that even with a human brain, other apes could not reach the human level of articulation and phonetic range^42, 44^. Phonetic range is determined by the different configurations that the vocal tract can produce. Modern humans have a 1:1 proportion between the horizontal and vertical dimensions of the vocal tract, which develops mainly in post-infant years^45^ and is unique among primates (Fig. 6a)^42^. Although it is still debated whether this configuration is a prerequisite for speech^43^, it was nonetheless suggested to be optimal for speech^42, 46^. The 1:1 proportion was reached through retraction of the human face, together with the descent of the larynx, pulling the tongue with it, and suggesting that the two process are tightly linked^47, 48^. In this regard, the fact that the top five skeletal DMGs regulate both facial protrusion and the anatomy of the larynx suggests that these two processes might have been linked, though the interaction between the two is still to be determined, as their exact developmental pathways are beyond the scope of the current study. For an in-depth review of the anatomy of vocalization and speech, see ^42^.

**Figure 6.**
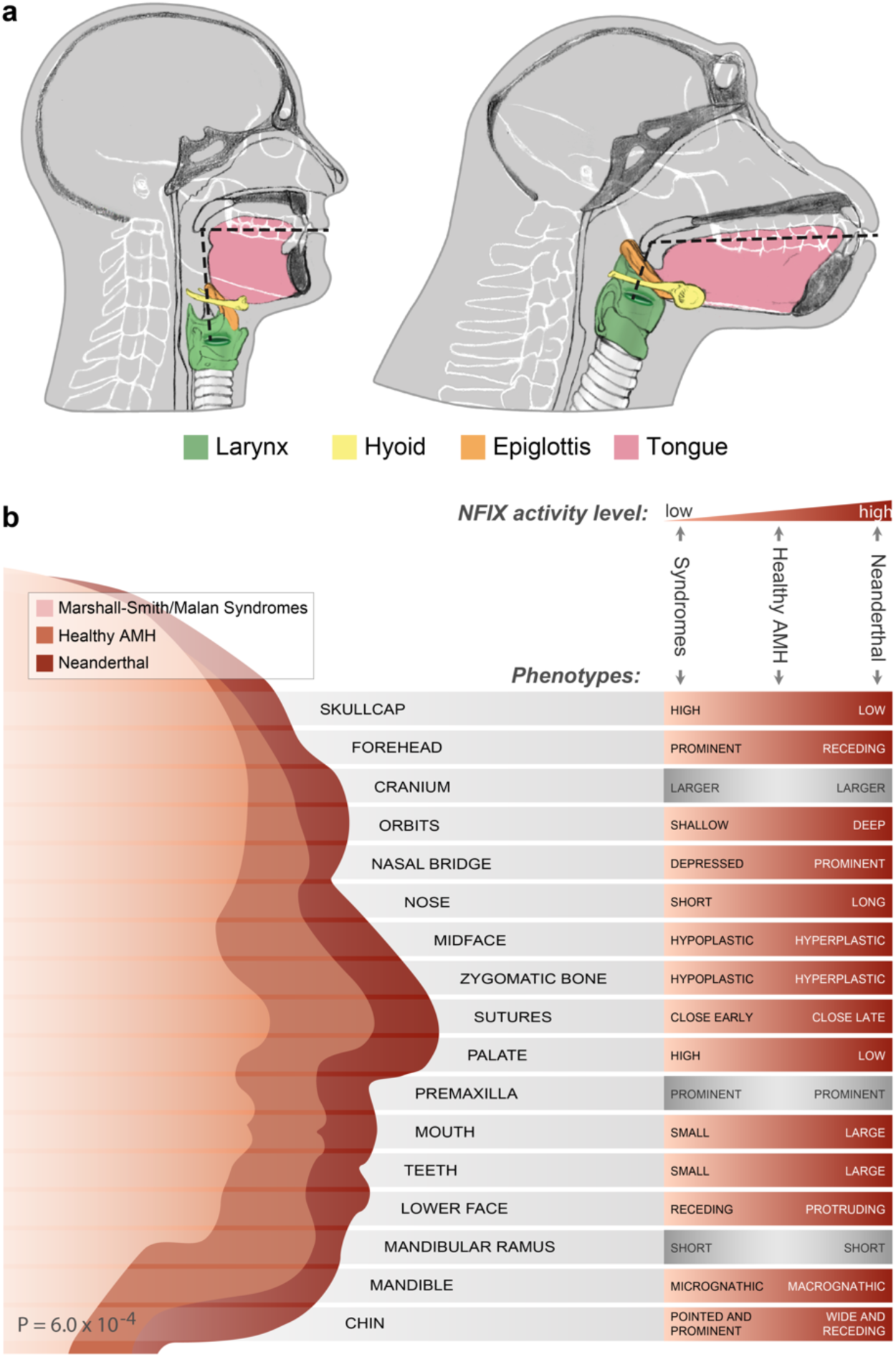
*NFIX* down-regulation may underlie modern human-derived traits. **a.** Vocal anatomy of chimpanzee and AMH. The vocal tract is the cavity from the lips to the larynx (marked by dashed lines). In AMHs, the flattening of the face together with the descent of the larynx led to approximately 1:1 proportions of the horizontal and vertical portions of the vocal tract. **b**. Craniofacial features of the Neanderthal, healthy AMH, and AMH with Marshall-Smith or Malan syndromes. Each box shows a phenotype that occurs in the Marshall-Smith/Malan syndromes (i.e., when *NFIX* is partially or completely inactive). The righthand side of each box shows the observed phenotypes of individuals with the syndromes (left), healthy AMHs (middle) and Neanderthals (right). In most phenotypes, the observed phenotypes match the expected phenotypes based on *NFIX* expression.

A longstanding question is whether Neanderthals and AMHs share similar vocal anatomy^49, 50^. Attempts to answer this question based on morphological differences have proven hard, as the larynx is mostly composed of soft tissues (e.g., cartilage), which do not survive long after death. The only remnant from the Neanderthal laryngeal region is the hyoid bone, which is detached from the rest of the skull^50^. Based on this single bone, or on computer simulations and tentative vocal tract reconstructions, it is difficult to characterize the full anatomy of the Neanderthal vocal apparatus, and opinions remain split as to whether it was similar to that of AMHs^49, 50^.

Most skeletal disease phenotypes that result from *NFIX* dysfunction are craniofacial, as *NFIX* influences the balance between lower and upper projection of the face^51^. In addition, mutations in *NFIX* were shown to impair speech capabilities^41, 52^. The exact mechanism is still unknown, but is thought to occur partly through skeletal alterations to the larynx^41^. To investigate if changes in *NFIX* expression could explain morphological changes in the AMH face and larynx, we examined its clinical skeletal phenotypes. Mutations in *NFIX* were shown to cause the Marshall-Smith and Malan syndromes, whose phenotypes include various skeletal alterations such as hypoplasia of the midface, retracted lower jaw, and depressed nasal bridge^51^. In many patients, the phenotypic alterations are driven by heterozygous loss-of-function mutations that cause haploinsufficiency. This shows that reduced activity of *NFIX*, even if partial, results in skeletal alterations^51^. Because *NFIX* is inferred to have been down-regulated in AMHs compared to archaic humans, we hypothesized that similar phenotypes to the ones that are driven by *NFIX* loss-of-function may also exist between modern and archaic humans. For example, because reduced activity of *NFIX* results in a more retracted face, we hypothesized that AMHs would present a more retracted face compared to archaic humans. We therefore examined the phenotypes of the Marshall-Smith and Malan syndromes and found that not only do most of these phenotypes exist between Neanderthals and modern humans, but their direction matches the direction expected from NFIX down-regulation along the AMH lineage (18 out of the 22 Marshall-Smith phenotypes, and 8 out of the 9 Malan phenotypes, *P* = 6.0 x 10^-4^, binomial test). In other words, from the Neanderthal, where *NFIX* activity is expected to be highest, through healthy AMHs, to individuals with *NFIX* haploinsufficiency, phenotypic manifestation matches the level of *NFIX* activity (Fig. 6b, Extended Data Table 8).

Notably, many cases of laryngeal malformations in the Marshall-Smith syndrome have been reported. Some of the patients exhibit positional changes to the larynx, changes in its width, and, more rarely, structural alterations to the arytenoid cartilage – the anchor point of the vocal folds, which controls their movement^53^. In fact, these laryngeal and facial changes are thought to underlie some of the limited speech capabilities observed in various patients^41^. This raises the possibility that *NFIX* down-regulation in AMHs might have driven phenotypic changes in the larynx too.

*SOX9*, *ACAN*, *COL2A1*, *XYLT1*, and *NFIX* are active in early stages of osteochondrogenesis, making the observation of differential methylation in mature bones puzzling at first glance. This could potentially be explained by two factors: (i) The methylome stabilizes as development progresses and remains largely unchanged from late development through adulthood. Thus, adult methylation states often reflect earlier development, and DMRs in adult stages often reflect DMRs in earlier activity levels^3,17, 54^. Therefore, these DMRs might reflect early methylation changes in mesenchymal progenitors that are carried over to later stages of osteogenesis. Indeed, the methylation patterns of *NFIX*, *SOX9*, *ACAN*, and *COL2A1* were shown to be established in early stages of human development and remain stable throughout differentiation from mesenchymal stem cells to mature osteocytes^55^. It is further supported by the observation that osteoblasts and chondrocytes show almost identical methylation levels in these DMRs, and are all as hypermethylated as the adult bone methylation levels we report^56^. We have reconfirmed this result by measuring methylation in these DMGs in primary human chondrocytes. Finally, we show that the upstream mesenchymal enhancer of *SOX9*[^23^] is differentially methylated in AMHs (Fig. 4b). (ii) Although expression levels of *SOX9*, *ACAN*, and *COL2A1* gradually decrease with skeletal maturation, these genes were shown to remain active in later developmental stages in the larynx, vertebrae, limbs, and jaws, including in their osteoblasts^21, 57^. Interestingly, these are also the organs that are most affected by mutations in these genes, implying that their late stages of activity might still play important roles in morphological patterning^20, 38–40^. It was also shown that facial growth patterns, which shape facial prognathism, differ between archaic and modern humans not only during early development, but also as late as adolescence^58^. Moreover, the main differences between human and chimp vocal tracts are established during post-infant years^45^. Although the DMRs we report most likely exist throughout the skeleton, including the larynx, the evidence we present for the cranium is more direct, as the patterns are observed in modern human and chimpanzee crania. Importantly, it has been suggested that the 1:1 vocal conformation could have been entirely driven by cranial, rather than laryngeal, alterations^48^. Once archaic human cranial samples are sequenced, these observations could be more directly tested.

The results we presented open a window to study the evolution of the human vocal tract and face from genetic and epigenetic perspectives. Our data suggest shared genetic mechanisms that shaped these anatomical regions and point to evolutionary events that separate AMHs from the Neanderthal and Denisovan. The mechanisms leading to such extensive regulatory shifts, as well as if and to what extent these evolutionary changes affected vocalization and speech capabilities are still to be determined.

## Acknowledgements

We would like to thank Sagiv Shifman, Yoel Rak, Rodrigo Lacruz, Erella Hovers, Anna Belfer-Cohen, Achinoam Blau, Iain Mathieson, Philip Lieberman, Daniel Lieberman, and Terry Capellini for their useful advice, Svante Pääbo, Janet Kelso, Kay Prüfer, Johannes Krause, and Anne Pusey for providing data, Sjaak Kaandorp and Christine Kaandorp-Huber from Safari park Beekse Bergen in Netherlands for their cooperation in animal conservation and use for research, and Maayan Harel for illustrations. D.G. is supported by the Clore Israel Foundation. TMB is supported by BFU2017-86471-P (MINECO/FEDER, UE), U01 MH106874 grant, Howard Hughes International Early Career, Obra Social “La Caixa” and Secretaria d’Universitats i Recerca and CERCA Programme del Departament d’Economia i Coneixement de la Generalitat de Catalunya. D.R. is an Investigator of the Howard Hughes Medical Institute and is also supported by an Allen Discovery Center for the Study of Human Brain Evolution funded the Paul G. Allen Family Foundation. C.L.-F. is supported by FEDER and BFU2015-64699-P grant from the Spanish government. R.P. was supported by ERC starting grant ADNABIOARC (263441). R.M.G. and J.M.O are supported by NYSTEM contract C030133. Funding for the collection and processing of the 850K chimp data was provided by the Leakey Foundation Research Grant for Doctoral Students, Wenner-Gren Foundation Dissertation Fieldwork Grant (Gr. 9310), James F. Nacey Fellowship from the Nacey Maggioncalda Foundation, International Primatological Society Research Grant, Sigma Xi Grant-in-Aid of Research, Center for Evolution and Medicine Venture Fund (ASU), Graduate Research and Support Program Grant (GPSA, ASU), and Graduate Student Research Grant (SHESC, ASU) to G.H. Collection of the chimpanzee bone from Tanzania was funded by the Jane Goodall Institute, and grants from the US National Institutes of Health (AI 058715) and National Science Foundation (IOS-1052693), and facilitated by Elizabeth Lonsdorf and Beatrice Hahn.

## Author Contributions

D.G. planned and conducted analyses. L.C. supervised the computational and experimental work. E.M. supervised experiments. L.A.T, B.Y, D.G. and L.C. conceived statistical analyses. D.G., L.C, and E.M. wrote the manuscript. All other authors contributed to the production of data and wrote their respective parts of the manuscript.

## Methods

### Skeletal Methylation Maps

Previously, our ability to identify differentially methylated regions (DMRs) that discriminate between human groups was confined by three main factors: (i) We had a single DNA methylation map from a present-day human bone, which was produced using a reduced representation bisulfite sequencing (RRBS) protocol, which provides information for only ∼10% of CpG positions in the genome. Moreover, the fact that the archaic and present-day methylomes were produced using different technologies – computational reconstruction versus RRBS – potentially introduces a bias. (ii) The analyses included only one bone methylation map from each of the human groups, which limited our ability to identify fixed differences between the groups. Although dozens of maps from additional tissues in present-day humans were included in the analyses, this narrowed the DMRs to represent only human-specific changes that are invariable between tissues. (iii) The work did not include a great ape outgroup. Thus, when a AMH-specific change was identified, it was impossible to determine whether it happened on the AMH lineage, or in the ancestor of Neanderthals and Denisovans^4^.

To overcome these obstacles, a major goal of the current study was to significantly extend the span of our skeletal methylome collection, covering as many individuals, sexes, and bone types as we could. This included the generation of many new samples, including the high-coverage sequencing of additional ancient genomes, as listed below.

#### Present-day human bone DNA methylation maps

We generated full DNA methylation maps from two femur head bones from present-day humans using whole-genome bisulfite sequencing (WGBS). Femora were chosen because of their abundance in present-day human samples, as well as in ancient DNA samples^5, 6, 59^. In addition, we collected 53 publicly available partial skeletal methylation maps.

##### WGBS of two modern human bones

###### Sample collection

Trabecular bone tissue from femur heads were taken from two patients with osteoarthritis during a total hip replacement surgery, and after filling in a consent form as per Helsinki approval #0178-13-HMO. Importantly, the effects of osteoarthritis processes on trabecular bone are much less substantial than those on the synovium, cartilage, and subchondral bone. Bone1 was a left head of femur taken on August 11, 2014 from a 66 years old female and Bone 2 was a right head of femur taken on September 2, 2014 from a 63 years old female.

###### DNA Extraction

DNA was extracted from bones using QIAamp® DNA Investigator kit (56504, Qiagen). Bones were cut to thin slices (0.2-0.5 mm) and then thoroughly washed (X5) with PBS, to clean samples from blood. Bones were crushed with mortar and pestle in liquid nitrogen, and 100 mg bone powder was taken to extract DNA according to the protocol “Isolation of Total DNA from Bones and Teeth” of the DNA Investigator kit.

###### WGBS

Whole-genome bisulfite sequencing was conducted at the Centre Nacional d’analysis Genomico (CNAG) as described in^60^. After cell sorting, genomic DNA libraries were constructed using the Illumina TruSeq Sample Preparation kit (Illumina) following the manufacturer’s standard protocol. DNA was then exposed to two rounds of sodium bisulfite treatment using the EpiTect Bisulfite kit (QIAGEN), and paired-end DNA sequencing was performed using the Illumina Hi-Seq 2000. We used the GEM mapper^61^ with two modified versions each of the human (GRCh37) and viral reference genomes: one with all C’s changed to T’s and another with all G’s changed to A’s. Reads were fully converted in silico prior to mapping to the modified reference genomes, and the original reads were restored after mapping. Although methylation state should not depend on read position, positional biases have been previously reported^62^. We observed that the first few bases from each read showed a slightly higher probability of being called as methylated, and we thus trimmed the first ten bases from each read (M-bias filtering).. Heterozygous positions, positions with a genotype error probability greater than 0.01, and positions with a read depth greater than 250 were filtered out. Only cytosines with six or more reads informative for methylation status were considered. On average, half of the reads from either strand will be informative for methylation status at a given position, so minimum coverage is typically greater than 12. Methylated and unmethylated cytosine conversion rates were determined from spiked-in bacteriophage DNA (fully methylated phage T7 and unmethylated phage lambda). Five samples were excluded based on conversion rates <0.997, supported by visual inspection of CG and non-CG methylation plots. The over-conversion rates for all samples based on methylated phage T7 DNA were ∼5%.

Sequence quality was evaluated using FastQC software v0.11.2. TRIMMOMATIC v.0-32 was used to filter low quality bases with the following parameters: -phred33 LEADING:30 TRAILING:30 MAXINFO:70:0.9 MINLEN:70. Paired-end sequencing reads were mapped to bisulfite converted human (hg19) reference genome using Bismark v0.14.3 and bowtie2 v2.2.4 not allowing multiple alignments and using the following parameters: --bowtie2 --non_bs_mm -- old_flag -p 4. Potential PCR duplicates were removed using Bismark’s deduplicate_bismark_alignment_output.pl Perl program. Bismark’s bismark_methylation_extractor script was used to produce methylation calls with the following parameters: -p --no_overlap --comprehensive --merge_non_CpG --no_header --bedGraph -- multicore 2 --cytosine_report. Examination of the M-bias plots led us to ignore the first 5 bp of both reads in human samples (Extended Data Fig. 5). Custom scripts were used to summarize methylation levels at CpG sites based on the frequencies of methylated and unmethylated mapped reads on both strands. Methylation data were deposited in NCBI’s Gene Expression Omnibus and are accessible through GEO accession number <u>GSE96833</u>.

##### Partial skeletal and full non-skeletal DNA Methylation maps of modern humans and chimpanzees

Osteoblast RRBS map, extracted from the femur, tibia, and rib bones of a 6-year-old female (NHOst-Osteoblasts by Lonza Pharma, product code: CC-2538, lot number: 6F4124), was downloaded from GEO accession number GSE27584. 48 450K methylation array maps, extracted from the femora of adult males and females with osteoarthritis or osteoporosis, were downloaded from GEO accession number GSE64490. Four 450K methylation array maps, extracted from unspecified bones of adult males and females were downloaded from GEO accession number GSE50192. Chimpanzee and human WGBS blood methylation maps were downloaded from NCBI SRA accession number SRP059313. Chimpanzee and human WGBS brain maps were downloaded from GEO accession number GSE37202.

##### Bisulfite-PCR of human bone

###### Sample collection

A skull of an adult male from India was obtained from the teaching anatomy collection of the Department of Anatomy and Anthropology at the Sackler Faculty of Medicine, Tel Aviv University, Israel (Human 1). Additional two skull specimens (Human 2 and 3) were obtained directly from the operating room of the Department of Neurosurgery, Shaare Zedek Medical Center, Jerusalem, Israel and transferred on dry ice for further analysis. All study participants provided informed consent according to an institutional review board – approved protocol (SZMC 0048-18).

###### DNA extraction

Human 1: Standard precautions to avoid contamination were taken, including wearing disposable coats, masks, hair covers and double gloves. All following steps were performed in a UV cabinet dedicated for the preparation of ancient bone samples and located in a physically separated ancient DNA laboratory at the Faculty of Dental Medicine. The skull was cleaned with an excess of 10% bleach (equal to 0.6% Sodium hypochlorite) and then subjected to UV radiation for 30 minutes. The cortical layer on the temporal surface (*facies temporalis*) of the zygomatic bone (ZB) was removed by low-speed drilling using a Wolf Multitool Combitool Rotary Multi Purpose Tool equipped with a sterile dental burr. Another sterile burr was used to obtain powder of the subcortical trabecular bone within the body of the zygoma. The powder was collected onto a 10 x 10 cm aluminum foil sheet pretreated with a 10% bleach solution and then transferred into a sterile 1.5 ml Eppendorf tube for subsequent DNA extraction. Altogether, three samples were obtained: ZB-3 from the right zygoma weighing 20.3 mg, and ZB-3/1 and ZB-3/2 from the left zygoma weighing 29.5 mg and 30.3 mg, respectively. Bone DNA was purified from the three bone powder samples using QIAamp DNA Investigator Kit (QIAgen, 56504) according to manufacturer’s instructions.

Human 2 and 3: DNA was extracted from bones using QIAamp® DNA Investigator kit (56504, Qiagen). Bones were thoroughly washed (X5) with PBS, to clean samples from blood. Bones were crushed with mortar and pestle in liquid nitrogen, and 100 mg bone powder was taken to extract DNA according to the protocol “Isolation of Total DNA from Bones and Teeth” of the DNA Investigator kit.

###### Bisulfite-PCR

Genomic DNA was bisulfite converted with the EZ DNA Methylation – Lightning Kit (Zymo Research, D5030) according to the manufacturer’s instructions. Specifically, each bone sample was bisulfite converted using 500ng as genomic DNA input for the conversion. Bisulfite treated DNA were amplified with the FastStart High Fidelity PCR System (Sigma, 03553400001) using the primers listed in Extended Data Table 12. PCR conditions were performed according to manufacturer’s instructions and PCR products were visualized on a 1.5 % agarose gel. Prior to cloning, PCR products were purified with Gel/PCR DNA Mini Kit (RBC, YDF100) and quantified with a NanoDrop 2000 spectrophotometer.

###### Cloning and sequencing

CloneJET PCR Cloning Kit (Thermo Scientific, K1231) was used to clone the purified PCR products into a pJET1.2/blunt Cloning Vector following the Blunt-End Cloning Protocol described in the manufacturer’s instructions. 5µl of each cloning reaction product were used for transformation of DH5α Competent Cells (Invitrogen, 18265017). Colonies were grown overnight on LB plates containing 100μg/ml ampicillin. Positive transformants were picked and grown overnight in liquid LB medium containing 100 μg/ml ampicillin. Subsequently, plasmid minipreps were purified with a RBC Miniprep Kit (YPD100) according to manufacturer’s instructions. Purified plasmids were quantified with a NanoDrop 2000 spectrophotometer and sequenced on an Applied Biosystems 3730xl Genetic Analyzer (Extended Data Fig. 3a,b).

###### Human primary chondrocyte validation

Primary chondrocyte cultures were obtained from osteoarthritis (OA) donors in accordance with Hadassah Medical Center Institutional Review Board approval and in accordance with the Helsinki Declaration of ethical principles for medical research involving human subjects. End-stage OA patients, with a Kellgren and Lawrence OA severity score of 3-4 were recruited following receipt of a formal written informed consent (n=8; 75% female, mean age 73±7.2 years; mean body mass index 30.1 ±5.4 kg/m^2^). Hyaline articular cartilage was dissected and human chondrocytes isolated using 3 mg/mL Collagenase Type II (Worthington Cat # LS004177) in DMEM medium (Sigma-Aldrich, St Louis, MI) containing 10% FCS and 1% penicillin-streptomycin (Beit-Haemek Kibutz, Israel), 37°C, 24h incubation. Isolated cells were filtered through a nylon cell strainer (40mm diameter), washed three times with PBS and plated at 1.5 million cells per 14 cm^2^ tissue culture dish (passage 0, passage 2). Cells were cultured in standard incubation conditions (37°C, 5% CO_2_) until confluence. Chondrocyte DNA purification was performed using GenElute™ Mammalian Genomic DNA Miniprep Kit (Sigma, G1N350).

#### Chimpanzee bone DNA methylation maps

Overall, we produced six methylation maps from bones of six common chimpanzee (*Pan troglodytes*) individuals. They include one WGBS of a wild chimpanzee, one RRBS of an infant chimpanzee, and four 850K methylation arrays of captive chimpanzees.

##### Ethics Statement

Chimpanzee tissue samples included in this study were opportunistically collected at routine necropsy of these animals. No animals were sacrificed for this study, and no living animals were used in this study.

##### WGBS of a chimpanzee bone

###### Sample collection

We used a rib bone of a 47-year-old female Chimpanzee provided from the Biobank of the Biomedical Primate Research Centre (BPRC), The Netherlands. The postmortem interval was approximately 10-12 hours. The bone was collected during the necropsy procedure and immediately frozen and stored at –80 ⁰C.

###### DNA extraction

DNA was extracted in a dedicated ancient DNA laboratory at the Institute of Evolutionary Biology in Barcelona, where no previous work on great apes has ever been conducted. Standard precautions to avoid and monitor exogenous contamination such as frequent cleaning of bench surfaces with bleach, use of sterile coveralls, UV irradiation and blank controls were taken during the process. 200 mg of bone powder were obtained by drilling and the sample was extracted following the Dabney et al. (2013) method^63^. A final 25 µL of extract volume was used for genome sequencing.

###### WGBS

Analysis was performed similarly to Bone1 and Bone2, with the exception that the BSreads were mapped to bisulfite converted chimpanzee (panTro4) reference genome, and we ignored the first 5bp of read1 and the first 44 bp of read2 in the chimpanzee sample (Extended Data Fig. 6).

Methylation data were deposited in NCBI’s Gene Expression Omnibus and are accessible through GEO accession number <u>GSE96833</u>.

##### RRBS of a chimpanzee bone

###### Sample collection

We used two unidentified long bone fragments that belonged to a newborn wild chimpanzee infant who died during a documented infanticide event at Gombe National Park on 9 March 2012. The infant was known to be the offspring of a chimpanzee called Eliza and was partially eaten by an adult female and her family. The sample was collected from the ground about 48 hours after the infant’s death and stored in RNAlater solution until arrival at Arizona State University (ASU). At ASU the sample was stored at 4°C until extraction.

###### DNA Extraction

Sampling and DNA extractions were conducted at the ASU Ancient DNA Laboratory, a Class 10,000 clean-room facility in a separate building from the Molecular Anthropology Laboratory. Precautions taken to avoid contamination included bleach decontamination and UV irradiation of tools and work area before and between uses, and use of full body coverings for all researchers. The bone samples were pulverized together in December 2012 using a SPEX CertiPrep Freezer Mill. Three DNA extractions were conducted using 50-100 mg of bone powder (Extended Data Table 9) and following the extraction protocol by Rohland and Hofreiter^64^. Two extraction blank controls were included to monitor contamination of the extraction process. One µL each of the sample extract and the blank control were used for fluorometric quantification with the Qubit 2.0 Broad Range assay. All extracts were combined for a total volume of 345 µL and approximately 0.652 µg of total DNA.

###### RRBS

RRBS libraries were generated according to Boyle *et al.*^65^. 100-200 ng genomic DNA was digested with MspI. Subsequently, the digested DNA fragments were end-repaired and adenylated in the same reaction. After ligation with methylated adapters, samples with different adapters were pooled together and were subjected to bisulfite conversion using the EpiTect Bisulfite kit (QIAGen) per the manufacturer’s recommendations with the following modification: after first bisulfite conversion, the converted DNA was treated with sodium bisulfite again to guarantee that conversion rates were no less than 99%. Two third of bisulfite converted DNA was PCR amplified and final RRBS libraries were sequenced in an Illumina HiSeq 2000 sequencer (Extended Data Table 10). Methylation data were deposited in NCBI’s Gene Expression Omnibus and are accessible through GEO accession number GSE96833.

##### 850K DNA methylation arrays

###### Sample collection

Four chimpanzee cadavers from captive colonies at the Southwest National Primate Research Center in Texas were used. Femora were opportunistically collected at routine necropsy of these animals and stored in −20°C freezers at the Texas Biomedical Research Institute after dissection. These preparation and storage conditions ensured the preservation of skeletal DNA methylation patterns.

###### DNA extraction

Samples were then transported to ASU and DNA was extracted from the femoral trabecular bone using a phenol-chloroform protocol optimized for skeletal tissues ^66^. From the distal femoral condyles, trabecular bone was collected using coring devices and pulverized into bone dust using a SPEX SamplePrep Freezer/Mill. Specifically, bone cores were obtained from a transverse plane through the center of the medial condyle on the right distal femur, such that the articular surface remained preserved. Cortical bone was removed from these cores using a Dremel (Extended Data Table 11). Tissue collections were performed at the Texas Biomedical Research Institute, and DNA extractions were conducted at the ASU Molecular Anthropology Laboratory.

###### Genome-Wide DNA Methylation Profiling

Genome-wide DNA methylation was assessed using Illumina Infinium MethylationEPIC microarrays. These arrays analyze the methylation status of over 850,000 sites throughout the genome, covering over 90% of the sites on the Infinium HumanMethylation450 BeadChip as well as an additional 350,000 sites within enhancer regions. For each sample, 400 ng of genomic DNA was bisulfite converted using the EZ DNA Methylation^TM^ Gold Kit according to the manufacturer’s instructions (Zymo Research), with modifications described in the Infinium Methylation Assay Protocol. These protocols were conducted at the ASU Molecular Anthropology Laboratory. Following manufacturer guidelines (Illumina), this processed DNA was then whole-genome amplified, enzymatically fragmented, hybridized to the arrays, and imaged using the Illumina iScan system. These protocols were conducted at the Texas Biomedical Research Institute. These array data have been deposited in NCBI’s Gene Expression Omnibus and are accessible through GEO Series accession number GSE94677.

###### Methylation Data Processing

Raw fluorescent data were normalized to account for the noise inherent within and between the arrays themselves. Specifically, we performed a normal-exponential out-of-band (Noob) background correction method with dye-bias normalization to adjust for background fluorescence and dye-based biases and followed this with a between-array normalization method (functional normalization) which removes unwanted variation by regressing out variability explained by the control probes present on the array as implemented in the minfi package in R ^67^ which is part of the Bioconductor project. This method has been found to outperform other existing approaches for studies that compare conditions with known large-scale differences ^67^, such as those assessed in this study.

After normalization, methylation values (β values) for each site were calculated as the ratio of methylated probe signal intensity to the sum of both methylated and unmethylated probe signal intensities. These β values range from 0 to 1 and represent the average methylation levels at each site across the entire population of cells from which DNA was extracted (0 = completely unmethylated sites, 1 = fully methylated sites).

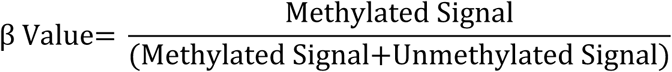

Every β value in the Infinium platform is accompanied by a detection p-value, and those with failed detection levels (p-value > 0.05) in greater than 10% of samples were removed from downstream analyses.

The probes on the arrays were designed to specifically hybridize with human DNA, so our use of chimpanzee DNA required that probes non-specific to the chimpanzee genome, which could produce biased methylation measurements, be computationally filtered out and excluded from downstream analyses. This was accomplished using methods modified from ^68^. Briefly, we used blastn to map the 866,837 50bp probes onto the chimpanzee genome (Assembly: Pan_tro_3.0, Accession: GCF_000001515.7) using an e-value threshold of e^-10^. We only retained probes that successfully mapped to the genome, had only 1 unique BLAST hit, targeted CpG sites, had 0 mismatches in 5bp closest to and including the CpG site, and had 0-2 mismatches in 45bp not including the CpG site. This filtering retained 622,819 probes.

Additionally, β values associated with cross-reactive probes, probes containing SNPs at the CpG site (either human or chimp), probes detecting SNP information, probes detecting methylation at non-CpG sites, and probes targeting sites within the sex chromosomes were removed using the minfi package in R ^67^. This filtering retained a final set of 576,505 probes.

##### Bisulfite-PCR of chimpanzee cranial bones

###### Sample collection

Postmortem frontal skull bones from two different chimpanzees (chimpanzee 1 and chimpanzee 2) were provided by the Biomedical Primate Research Centre (BPRC, The Netherlands). Bones were opportunistically collected during routine necropsy of these animals and stored at −80°C. Chimpanzee 3 and chimpanzee 4 samples were obtained from the chimpanzee cranial collection in the Department of Paleoanthropology in the Senckenberg Research Institute Frankfurt (DPSF) and Natural History Museum Frankfurt ^69^. These two chimpanzee specimens are owned by the Justus Liebig University Gießen.

###### DNA extraction

*Chimpanzee 1 and chimpanzee 2*

For each sample, bone powder was obtained by crushing the bones with mortar and pestle. Approximately 100mg bone powder were used to extract DNA using the QIAamp DNA Investigator Kit (Qiagen) following manufacturer’s instructions.

Chimpanzee 3 and chimpanzee 4

Cochlear bone powder was obtained by accessing the petrous bone from the cranial base^70^. DNA was extracted from about 50 mg of powder according to the protocol described by^63^, but adapted for the use of High Pure Nucleic Acid Large Volume columns (Roche) instead of the Zymo-Spin V column (Zymo Research) MinElute silica spin column (Qiagen) combination.

###### Bisulfite-PCR

Genomic DNA was bisulfite converted with the EZ DNA Methylation – Lightning Kit (Zymo Research, D5030) according to the manufacturer’s instructions. Specifically, each bone sample was bisulfite converted two times in parallel using 500ng as genomic DNA input for the conversion.

3µl of bisulfite treated DNA were amplified with the FastStart High Fidelity PCR System (Sigma, 03553400001) using the primers listed in Extended Data Table 12. PCR conditions were performed according to manufacturer’s instructions and PCR products were visualized on a 1.5 % agarose gel. Prior to cloning, PCR products were purified with homemade SPRI beads (chimpanzee 1 and 2) and Gel/PCR DNA Mini Kit (RBC, YDF100, chimpanzee 3 and 4), and quantified with a NanoDrop 2000 spectrophotometer.

###### Cloning and sequencing

CloneJET PCR Cloning Kit (Thermo Scientific, K1231) was used to clone the purified PCR products into a pJET1.2/blunt Cloning Vector following the Blunt-End Cloning Protocol described in the manufacturer’s instructions. 3µl (chimpanzee 1 and 2) and 3µl (chimpanzee 3 and 4) of each cloning reaction product were used for transformation of DH5α Competent Cells (Invitrogen, 18265017). Colonies were grown overnight on LB plates containing 100μg/ml ampicillin. Positive transformants were picked and grown overnight in liquid LB medium containing 100 μg/ml ampicillin. Subsequently, plasmid minipreps were purified with a QIAprep Miniprep Kit (Qiagen, chimpanzee 1 and 2), and RBC Miniprep Kit (YPD100, chimpanzee 3 and 4) according to manufacturer’s instructions. Purified plasmids were quantified with a NanoDrop 2000 spectrophotometer and sequenced on an Applied Biosystems 3730xl Genetic Analyzer (Extended Data Fig. 3a,b).

### Reconstructing ancient DNA methylation maps

#### La Braña 1 genome sequencing

In a dedicated clean room at Harvard Medical School, powder was extracted from the root of a lower third molar of the Mesolithic La Braña 1 individual (5983-5747 calBCE (6980±50 BP, Beta-226472)), from which a non-UDG-treated library was previously sequenced to 3.5x coverage^8^. Two UDG-treated libraries from the same individual were later generated and enriched for approximately 1.2 million single targeted polymorphisms and sequenced to an average of 19.5x coverage at these positions ^9^. In this study, we carried out shotgun sequencing of one of the same UDG-treated libraries from this individual on a NextSeq500 instrument using 2 x 76bp paired end sequences ^71^. Following the mapping protocol described previously ^9^, we trimmed adapter sequences, only processed read pairs whose ends overlapped by at least 15 bp (allowing for one mismatch) so that we could confidently merge them, and then mapped to the human reference sequence hg19 using the command samse in BWA (v0.6.1). We removed duplicated sequences by identifying sequences with the same start and stop position and orientation in the alignment, and picking the highest quality one. After restricting to sequences with a map quality of MAPQ ≥ 10, and sites with a minimum sequencing quality (≥20), we had an average coverage measured at the same set of approximately 1.2 million single nucleotide polymorphism targets of 23.0x. This data is available under GEO accession number: GSE96833, with raw reads deposited under SRA accession number: SRX3194436.

#### I1583 Genome sequencing

In a dedicated clean room at the University College Dublin, powder was extracted from the cochlear portion of the petrous bone of individual I1583 (archaeological ID L14-200) from the site of Barcın Höyük in the Yenişehir Plain of the Marmara Region of Northwest Turkey. The Neolithic individual came from a community that practiced farming, and was anthropologically determined to be a male aged 6-10 years at the time of death (the sex was confirmed genetically). The direct radiocarbon date was 6426-6236 calBCE (7460±50 BP, Poz-82231). In a dedicated clean room at Harvard Medical School, a UDG-treated library was prepared from this powder, which was previously enriched for about 1.2 million SNP targets, sequenced to 13.5x average coverage, and published in ^9^. We shotgun sequenced the same library on nine lanes of a HiSeqX10 sequencing with 100bp paired reads. On data processing, we merged overlapping read pairs, trimmed Illumina sequencing adapters, and dropped read pairs that did not have sample barcodes (up to 1 mismatch) or cannot be unambiguously merged. We then aligned merged reads with BWA against human reference genome GRCh37 (hg19) plus decoy sequences, and combined all nine lanes of data and removed duplicate molecules, achieving an average of 24.3x coverage evaluated on the 1.2 million targets. This data is available under GEO accession number: GSE96833, with raw reads deposited under SRA accession number: SRX3194436.

#### The reconstruction procedure

Reconstruction of DNA methylation maps was performed on the genomes of the following individuals: Ust’-Ishim^6^, Loschbour^7^, Stuttgart^7^, La Braña 1, I1583, and the Vindija Neanderthal^5^, as well as on the previously published Altai Neanderthal and the Denisovan (Extended Data Table 1). The Vindija Neanderthal reads were downloaded from the Max Planck Institute for Evolutionary Anthropology website: http://cdna.eva.mpg.de/neandertal/Vindija/bam/. Only the UDG-treated portion of the genome (B8744) was used. Additional UDG-treated ancient human full genomes have been published to date; however, these were sequenced to a relatively low coverage (<5x), and thus, only crude methylation maps could be reconstructed from them. C→T ratio was computed for every CpG position along the hg19 (GRCh37) human genome assembly, for each of the samples, as previously described^4^.

In order to exclude from the analyses positions that potentially represent pre-mortem C→T mutations rather than post-mortem deamination, the following filters were applied: (i) Positions where the sum of A and G reads was greater than the sum of C and T reads were excluded. (ii) For genomes that were produced using single-stranded libraries (i.e., Ust’-Ishim, Altai Neanderthal, Denisovan, Vindija Neanderthal and ∼1/3 of the Loschbour library), positions where the G→A ratio on the opposite strand was greater than 1/(average single strand coverage) were excluded. This fraction represents a threshold of one sequencing error allowed per position. For Loschbour, this was performed only on the fraction of reads that came from the single stranded library. (iii) For all genomes, positions with a C→T ratio > 0.25 were discarded. For the Vindija Neanderthal, this threshold was raised to 0.5, due to its relatively low coverage (∼7x). (iv) Finally, a maximum coverage threshold of 100 reads was used to filter out regions that are suspected to be PCR duplicates.

In all genomes, excluding Vindija, a fixed sliding window of 25 CpGs was used for smoothing of the C→T ratio. This allowed for an unbiased scanning of differentially methylated regions (DMRs) that is not affected by the size of the window. Due to its relatively low coverage, we extended the sliding window used on the Vindija genome to 50 CpGs. This extended window is not expected to introduce a bias, as this genome was not used for DMR detection, but only for subsequent filtering that was applied equally to all genomes (see later).

As previously described, C→T ratio was translated to methylation percentage using linear transformation determined from two points: zero C→T ratio was set to the value 0% methylation, and mean C→T ratio in completely methylated (100% methylation) CpG positions in modern human bone reference (hereinafter μ_100_) was set to the value 100% methylation. Positions where C→T ratio > μ_100_ were set to 100% methylation. For genomes that were extracted from bones, the modern Bone 2 WGBS map, which is the one with the higher coverage between the two WGBS modern bone maps, was used to determine μ_100_. For genomes that were extracted from teeth, there was no available modern reference methylation map, and therefore, we transformed the C→T ratio into methylation percentage based on the assumption that the genome-wide mean methylation is similar to bone tissue. Thus, the genome-wide mean C→T ratio represents 75% methylation, which is the genome-wide mean of measured methylation in the Bone 2 reference map. This was accomplished by setting μ_100_ to 1.33 x mean genome-wide C→T ratio.

### DMR detection

The DMR detection algorithm is comprised of five main steps. We hereby provide an overview of the algorithm followed by a detailed description of each step. The overall goal of this pipeline is to detect differential methylation, assign it to the lineage on which it arose and filter out within-lineage variation.

#### Overview

Step 1: Two-way comparisons. To avoid artifacts that could potentially be introduced by comparing DNA methylation maps that were produced using different technologies, our core analysis relied on the comparison of the three reconstructed maps of the Altai Neanderthal, Denisovan, and Ust’-Ishim. Each of the samples was compared to the other two in a pair-wise manner, as a raw C→T ratio map against a reconstructed methylation map, and vice versa. This reciprocal comparison insured that the reconstruction process does not introduce biases to one of the groups. The minimum methylation difference threshold was set to 50%, spanning >50 CpGs. Step 2: Three-way comparisons. This step classifies to which of the three hominins the DMR should be attributed. This step is done by overlapping the three lists of DMRs found in Step 1.

For example, a DMR that is detected between the Neanderthal and Ust’-Ishim and also between the Denisovan and Ust’-Ishim is considered specific to Ust’-Ishim.

Step 3: FDR filtering. Various factors could introduce noise to the reconstruction process, including the stochasticity of the deamination process, the use of a sliding window, and variations in read depth within a sample. We ran simulations that mimic the post-mortem degradation processes of ancient DNA, then reconstructed methylation maps from the simulated deamination maps and finally compared them to the original map and identified DMRs. Any differences in methylation levels between the simulated map and the original reference map stem from noise. Thus, running the same DMR-detection algorithm on the simulated map vs. the reference map, enables an estimation of the false discovery rate. We set the DMR-detection thresholds so that FDR < 0.05.

Step 4: Lineage assignment. The chimpanzee methylation maps were used to polarize the DMRs. For each DMR, methylation levels in the chimpanzee were compared to those of the three hominin groups. For example, if methylation levels in the chimpanzee samples clustered with the archaic humans, the DMR was assigned to the AMH lineage.

Step 5: Within-lineage variability filtering. To determine whether a DMR represents an individual within a group, or is shared by the entire group, we used a total of 67 AMH, archaic and chimpanzee methylation maps. We used a conservative approach where DMRs in which methylation levels in one group overlap (even partially) the methylation levels in another group were discarded. As 59 out of the 67 maps belong to AMHs, our ability to filter out variation within this group was better, resulting in fewer DMRs along this lineage. Several various measures were used to ascertain that a DMR along a lineage does not represent a sex-, bone-, age-, technology or disease-specific DMR.

#### DMR-detection algorithm

We developed an algorithm specifically designed to identify DMRs between a deamination map and a full methylome reference. Let *i* enumerate the CpG positions in the genome. In the deamination map, let *t_i_* be the number of T’s at the C position + the number of A’s in the opposite strand at the G position, i.e., it counts the total number of T’s that appear in a position that is originally C, in the context of a CpG dinucleotide. We similarly use *c_i_* to count the total number of C’s that appear in a position that is originally C, in the context of a CpG dinucleotide. The C→T ratio is defined as *t_i_*/*n_i_*, where *n_i_* = *c_i_* + *t_i_*. Let *φ_i_* and *ψ_i_* (both between zero and one) be the methylation of this position in the reference genome and in the reconstructed one, respectively. If we denote by **π** the deamination rate, assumed to be constant throughout the genome, and if we assume that deamination of C into T is a binomial process with probability of success *π*ψ_i_, we get

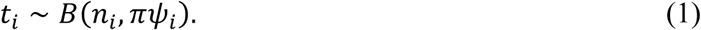

Our null hypothesis is that the *i*^th^ CpG is not part of a DMR, namely that *ψ_i_* = *φ_i_*. The alternative hypothesis states that this CpG is part of a DMR. The definition of this statement is that |*ψ_i_* − *φ_i_*| ≥ Δ, where Δ is some pre-specified threshold. In other words, under the alternative hypothesis we get that *ψ_i_* ≥ *φ_i_* + Δ if the site has low methylation in the reference genome, and *ψ_i_* ≤ *φ_i_* − Δ if it has high methylation in the reference genome.

##### Per-site statistic

Let us start with the first option, testing whether *ψ_i_* ≥ *φ_i_* + Δ when *φ_i_* is low. A log-likelihood-ratio statistic would be

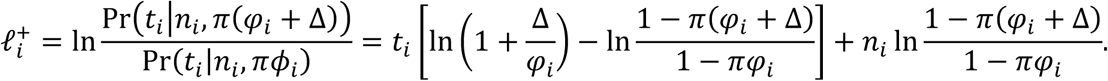

Similarly, we can test whether *ψ_i_* ≤ *φ_i_* − Δ when *φ_i_* is high using the log-likelihood-ratio statistic

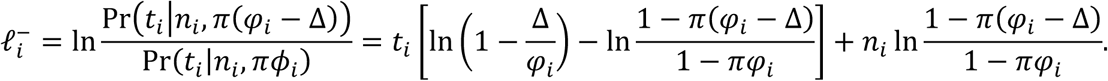

We used the value Δ = 0.5 for all samples. The value of *π*, the deamination rate, was estimated using the overall C→T ratio in CpG positions whose methylation level is 1 in the modern human Bone 2 WGBS methylation map, after exclusion of putative pre-mortem substitutions, as described in the “the reconstruction procedure” section (Extended Data Table 1).

##### Detecting DMRs

The statistics 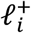 and 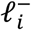 quantify how strongly the estimated methylation in position *i* deviates from *φ_i_*. Next, we use these values to identify DMRs using the cumulative-sum procedure explained below. The process is repeated twice: on the statistic 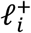 to identify DMRs where the sample has elevated methylation with respect to the reference, and on the statistic 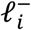 to identify DMRs where the sample has reduced methylation with respect to the reference.

For convenience, we explain the cumulative-sum procedure in the context of 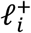, but an essentially identical procedure is used for 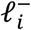. We define a new vector *Q*^+^ by the recursion

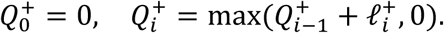

Under the null hypothesis, 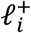 has a negative expectation which produces a negative drift that keeps *Q*^+^ at zero, or close to zero, levels. Under the alternative hypothesis the expectation is positive, hence the drift over a DMR is positive, leading to an elevation in the values of *Q*^+^. Therefore, our next step is to find all intervals [*a*, *b*] such that 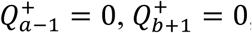, and 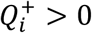 for *a* ≤ *i* ≤ *b*. Let 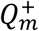 be the maximum value of *Q*^+^ in this interval, where *m* is the position of the maximum. Then, the interval [*a*, *m*] would be called a putative DMR.

The statistics 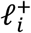 and 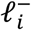 are affected by the number of observed cytosine reads, and thus have higher power to detect hypermethylation (i.e., larger number of cytosine reads) vs. hypomethylation (Extended Data Fig. 2).

##### Filtering DMRs

Of course, *Q*^+^ may increase locally due to randomness, and thus a putative DMR may not reflect a true DMR. To filter out such intervals, we used two strategies. First, we applied a set of filters to assure that the putative DMRs have reasonable biological properties. Second, we cleaned the remaining putative DMRs by applying a false discovery rate (FDR) procedure. In the first strategy, we applied two filters: (i) Putative DMRs that harbor less than 50 CpG positions, thus are shorter than twice the smoothing window size, were removed. (ii) To avoid situations where two consecutive CpG sites whose genomic locations are remote appear on the same DMR, we modify the vector 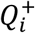 as follows. Let *d_i,j_* be the distance along the genome (in nucleotides) between CpG sites *i* and *j*. Then, for every site *i* such that *d_i,j-1_* > *δ* we set 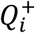 = 0. We used *δ* = 1000 nt for all samples.

To further remove putative DMRs that are unlikely to reflect true DMRs, we eliminated all DMRs where 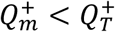. Here, 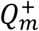 is the maximum value of *Q*^+^ in the interval as defined earlier, and 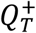 is a threshold determined using a false discovery rate (FDR) procedure, see section “filtering out noise” below.

##### Testing the algorithm

To verify that the approach above results in a low number of false positives, we applied the procedure to deamination maps, when compared to themselves in the form of reconstructed methylomes. As expected, we obtained a negligible number of DMRs, ranging between 0.4% and 1% of the number of DMRs detected between the humans.

#### Two-way DMR detection

In order to avoid artifacts that could potentially be introduced by comparing DNA methylation maps that were produced using different technologies, our core analysis relied on the comparison of the three reconstructed maps of the Altai Neanderthal, Denisovan, and Ust’-Ishim. These are all high-resolution maps that were derived from genomes sequenced to high coverage (Extended Data Table 1). In particular, the Ust’-Ishim methylome is of exceptional quality due to its high coverage and deamination rate (Extended Data Table 1). Also, going through the same post-mortem degradation processes, the Ust’-Ishim cellular composition is likely to be similar to that of the Neanderthal and Denisovan.

In order for a deamination map to serve as a reference in the comparison, we have transformed its C→T ratio values into methylation values (see “the reconstruction procedure” section above). To remove potential bias that could be introduced through the comparison of a reconstructed methylation map to a deamination map, we ran each two-way comparison twice: once with the methylation map of sample 1 against the deamination map of sample 2, and once with the deamination map of sample 1 against the methylation map of sample 2 (Extended Data Fig. 1). Therefore, the comparison of three genomes required a total of six two-way comparisons: Ust’-Ishim versus an Altai Neanderthal reference, Ust’-Ishim versus a Denisovan reference, Altai Neanderthal versus an Ust’-Ishim reference, Altai Neanderthal versus a Denisovan reference, Denisovan versus Ust’-Ishim reference, and Denisovan versus Altai Neanderthal reference. Because the DNA of these three individuals was extracted from both sexes, the DMR-detection algorithm was only applied to autosomes.

#### Three-way DMR detection

In order to identify DMRs where one group of humans (hereinafter, hominin 1) differs from the other two human groups (hereinafter, hominin 2 and hominin 3), we set out to find those DMRs that were detected both between hominin 1 and 2, and between hominin 1 and 3. To this end, we compare the two lists (hominin 1 vs. hominin 2 and hominin 1 vs. hominin 3) and look for overlapping DMRs, as previously described^4^. An overlapping DMR exists when a DMR from one list partially (or fully) overlaps a DMR from the second list. Only the overlapping portion of the two DMRs from the two-lists was taken.

#### Filtering out noise

There are different factors that potentially introduce noise into the reconstruction process. These include the stochasticity of the deamination process, the use of a sliding window to smooth the C→T signal, and variations in read depth. In order to account for these factors and estimate noise levels, we ran simulations that mimic the post-mortem degradation processes of ancient DNA, then reconstructed methylation maps from the simulated deamination maps and finally compared them to the original map and identified DMRs.

The simulation process starts with a methylation map, where the measured or reconstructed methylation at position *i* is *ψ_i_* and is assumed the true methylation. Given that *n_i_* is the coverage at this position, we use the binomial distribution (1) to randomly draw *t_i_* – the number of C’s that had become T’s through deamination. The resulting *t_i_*’s were then used to compute the C→T ratios for each position, smoothed and filtered using the same sliding window and thresholds used in the original analysis, and linearly transformed to methylation percentages as explained above (hereinafter, simulated methylation map, Extended Data Fig. 4a). Any differences in methylation levels between the simulated map and the original reference map stem from noise. Thus, running the same DMR-detection algorithm described above on the simulated map vs. the reference map, enables an estimation of the false discovery rate. We ran these simulations 100 times for each of the three genomes (Altai Neanderthal, Denisovan, Ust’-Ishim) and determined the values of the 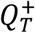 and 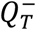 thresholds (see section “filtering DMRs” above) such that the mean number of DMRs that are detected in the simulations is < 0.05 the number of real DMRs detected (i.e., FDR < 0.05).

#### DMRs separating chimpanzees and humans

To identify DMRs that separate chimpanzees from all human groups (both modern and archaic), we first compared the chimpanzee WGBS bone methylome to each of two present-day WGBS maps (those of Bone1 and Bone2). This was done by scanning the chimpanzee map using a sliding window of 25 CpGs, in intervals of one CpG position. In each window, we counted the number of methylated and unmethylated reads in each sample, and computed a *P*-value using Fisher’s Exact test. We then computed FDR-adjusted *P*-values for each window, and discarded windows with FDR > 0.05 or where the mean methylation difference (Δ) was below 0.5. We then merged overlapping windows. This left 8,040 DMRs between the chimpanzee and Bone1, and 12,666 DMRs between the chimpanzee and Bone2. Next, we intersected the two lists to identify DMRs where the chimpanzee differs from the both present-day samples. This left 6,417 DMRs. Lastly, we compared the chimpanzee methylation levels to all other human samples (modern and archaic) and filtered out DMRs where the chimpanzee is found within the range of methylation levels observed in humans. To do so, we followed the procedure described in the “Removing DMRs with high within-group variability” section below. This resulted in 2,031 DMRs that separate chimpanzees and humans.

### Determining the lineages where DMRs originated

DMRs where Ust’-Ishim differs from the Neanderthal and the Denisovan could either arose on the AMH branch, or in the ancestor of Neanderthals and Denisovans. In order to allocate the DMRs to the branch in which the change occurred, we used the chimpanzee DNA methylation data.

First, we used the chimpanzee bone WGBS map. We defined the distance of a DMR in hominin *H* to chimpanzee as the mean absolute difference in methylation, 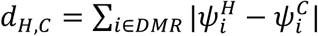. Here, 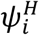 is the reconstructed methylation at the *i*’th CpG in hominin *H*, and 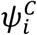 is the measured methylation in the same site in the chimpanzee. For Ust’-Ishim-specific DMRs, we used the following procedure: (i) If both archaic humans were closer to the chimpanzee, the DMR was placed on the AMH branch. (ii) If Ust’-Ishim was closer than both archaic humans to the chimpanzee, the DMR was placed on the branch of the ancestor of Neanderthals and Denisovans. (iii) Otherwise, the DMR was discarded. Out of 5,111 Ust’-Ishim-specific DMRs, we could place 1,729 DMRs on the AMH branch and 1,255 on the branch of the ancestor of Neanderthals and Denisovans. 1,807 Ust’-Ishim-specific DMRs were discarded due to inconclusive lineage assignment, and 320 had no data in the chimpanzee WGBS map. For Neanderthal-specific DMRs, we discarded all DMRs where Ust’-Ishim and the Denisovan were not found to be closer to the chimpanzee than the Neanderthal. Out of 3,107 Neanderthal-specific DMRs, 693 were placed on the Neanderthal branch, 2,202 were deemed inconclusive and were discarded, and 212 had no data in the chimpanzee WGBS map. Similarly, we discarded Denisovan-specific DMRs where Ust’-Ishim and Altai Neanderthal were not found to be closer to the chimpanzee than the Denisovan. Out of 1,461 Denisovan-specific DMRs, 499 were placed on the Denisovan branch, 855 were deemed inconclusive, and for 107 we had no data in the chimpanzee WGBS map. We next developed a second, stricter, scheme by also using the chimp 850K DNA methylation arrays datasets. As the probes cover just part of the CpGs in a DMR, we need to adjust the DMR methylation level in order to allow a meaningful comparison of 850K methylation data to full methylation maps. If we mark by *j* the CpGs in a DMR that are covered by 850K methylation array (which is a subset of all the CpGs in this DMR), and mark their total number by *J* = Σ_j∈DMR_ 1, then the methylation in the DMR as measured by the array is *m* = 1/*J*. 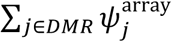, where 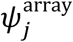 is the methylation level measured at position *j* in the array. Let 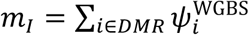 be the methylation of this DMR as computed from the full methylation map, where 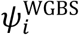 is the methylation level measured at position *i* in the full map. Let *m_J_* = 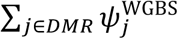 be the methylation as computed from the full methylation map when limited only to positions. Then, we correct the array methylation value *m* to:

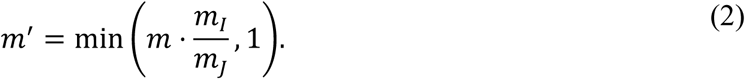

This procedure was applied to DMRs covered by at least one probe (∼65% of DMRs). For the remaining ∼35% of DMRs, we only used the WGBS chimpanzee methylome. This approach was used in parallel with filtering DMRs using the modern human 450K arrays (Extended Data Fig. 1, see next section).

There are pros and cons to each of these approaches. Using more chimpanzee datasets allow for more informative process. However, 850K methylation array probes are distributed unevenly across the genome. Although most DMRs are covered by at least one probe (mean number of probes per DMR: 1.7, median: 1, maximum: 64), many are nonetheless not covered. AMH On one hand, lineage assignment of DMRs for which we have array data is more robust and less prone to misclassification. On the other hand, DMRs with array data are more likely to be filtered out, as there is more power to detect variability. This could potentially alter the genomic distribution of DMRs. Therefore, we use both approaches throughout the paper. In analyses where it is important to maintain an unbiased distribution of DMRs we only use the chimpanzee WGBS map for polarization, and AMH bone WGBS maps for filtering (see next chapter), whereas in analyses where it is more important to minimize variability, or where we look at specific DMRs, we use the stricter approach. The chimpanzee RRBS data was adjusted using the same technique. However, it was not used for lineage assignment, but rather only as a source for additional information on DMRs. This is because this protocol particularly targets unmethylated CpGs, and is therefore too biased for lineage assignment.

### Removing DMRs with high within-group variability

Our three-way DMR detection algorithm above produces a list of DMRs where one of the three hominins (Ust’-Ishim, Altai Neanderthal or Denisovan) is significantly different from the other two. However, such DMRs could stem from variability within any of the groups, and in such cases cannot be regarded as truly differentiating between the human groups. Some variability may be removed during the process described above, in “Determining the lineages where DMRs originate”, but even DMRs whose origin can be assigned to a particular lineage do not necessarily represent fixed methylation changes. To filter out regions that are variable within any of the human groups, or across all of them, we used two approaches. First, we used the two modern human WGBS maps, and the I1583 reconstructed skull methylation map. DMRs where the Neanderthal or Denisovan methylation levels were found within the range of modern human methylation (i.e., Ust’-Ishim, the two WGBS maps and I1583) were discarded. This left 1,530 out of 1,729 Ust’-Ishim-derived DMRs (hereinafter, full AMH-derived DMRs), 1,230 out of 1,255 DMRs where the Neanderthal and Denisovan are both derived, 692 out of 693 full Neanderthal-derived DMRs, and 496 out of 499 Denisovan-derived DMRs.

The second approach adds to this the 52 450K methylation array samples, as well as the three reconstructed methylation maps from teeth (i.e., Loschbour, Stuttgart and La Braña 1). As described above, using also methylation probes for filtering DMRs provides more power, but can also introduce biases. Thus, this filtering was used for most analyses, except those where unbiased genomic distribution of DMRs is critical. Probe methylation data was corrected as described in equation (2). Within AMH- and archaic-derived DMRs, a DMR was deemed fixed if the Neanderthal and the Denisovan methylation levels both fell outside the range of methylation across all modern human samples (reconstructed, WGBS and 450K maps).

Similarly, within Neanderthal- and Denisovan-derived DMRs, a DMR was deemed fixed if the respective hominin fell outside the range of methylation across all modern human samples and the other archaic hominin. This approach yielded 873 AMH-derived DMRs (hereinafter referred to as AMH-derived DMRs), 939 archaic-derived DMRs, 570 Neanderthal-derived DMRs, and 443 Denisovan-derived DMRs.

The limited number of archaic human methylation maps introduces asymmetry in our ability to determine the level of fixation of DMRs along different lineages. Whereas we used dozens of AMH skeletal samples, we have just a few archaic samples. This provides us with the ability to better estimate the distribution of methylation values within each DMR in AMH, and thus to determine how significantly methylation values in other samples deviate from it. To enhance our ability to estimate variability within archaic human lineages, we added to the analysis the reconstructed methylation map of the Vindija Neanderthal. The USER-treated portion of this genome (the portion amenable for methylation reconstruction) was sequenced to a depth of 7x ^5^. Therefore, the methylation map that could be reconstructed from this individual has a considerably lower resolution compared to the other reconstructed maps used in this study (coverage 19x to 52x). Nevertheless, due to the reduced ability to detect variability along the archaic human linages, we employed this map for additional variability filtering along these lineages. DMRs where the Vindija Neanderthal clustered with the other hominins, and not with the Altai Neanderthal (or not with either of the archaic humans in the archaic-derived DMRs) were discarded. The number of DMRs mentioned throughout this chapter already includes this filtering.

A general concern in working with DNA methylation data is that DMRs that are specific to one group do not necessarily represent an evolutionary change, but rather reflect a characteristic such as technology used to measure methylation, tissue, sex, disease or age that is shared by individuals in this group and not by others. We take two complementary approaches to ascertain that the DMRs we report are not driven by these factors: (a) for the top DMGs, we match the samples for the above factors and test whether the hypermethylation of AMHs is still observed. To this end, we compared Ust’-Ishim (adult femur with no known diseases, methylation map produced using our reconstruction method) to the Vindija Neanderthal (adult femur with no known diseases, methylation map produced using our reconstruction method), and we also compared 52 modern human samples (adult femora, methylation array maps) to four chimpanzee samples (adult femora, methylation array maps). In all cases, AMHs show significant hypermethylation compared to the matched samples (Extended Data Figure 3c,d, see “*Comparing SOX9, ACAN, COL2A1, XYLT1, and NFIX methylation between AMH, chimpanzee and Neanderthal femora*” chapter for additional information). (b) throughout the pipeline, we take only DMRs where one human group clusters completely outside the other groups regardless of tissue, sex, disease, age or technology. Thus, these factors are unlikely to drive the reported methylation changes. This approach is particularly useful in AMH-derived DMRs, where each group of samples (i.e., AMH samples vs. archaic and chimpanzee samples) include both males and females, juveniles and adults, and they come from femora, ribs, tibia, skulls, and teeth. Thus, it is unlikely that the DMRs that differentiate these groups reflect variability that stems from these parameters^72^ (Fig. 1a-c). Archaic-derived DMRs and Neanderthal-derived DMRs are also unlikely to reflect differences in the above parameters, as in these DMRs, the Vindija Neanderthal sample (adult, femur bone) is clustered with the Altai Neanderthal sample (juvenile/adult, phalanx), and not with AMHs, where most samples are from femora of adult females. Denisovan-derived DMRs, on the other hand, are more likely to stem from age or bone type differences than other types of DMRs. This is because the Denisovan sample is the only finger bone, and it comes from a child (6-13.5 years) (Extended Data Table 1). Thus, we cannot rule out the possibility that some of the Denisovan-derived DMRs reflect finger-specific, rather than lineage-specific methylation patterns. These DMRs could also possibly reflect age-specific differences, but this is less likely, as the AMH I1583 sample and the chimpanzee 850K samples are the same age group as the Denisovan (Denisovan: 6-13.5 years old, I1583: 6-10 ^9^, chimpanzees: 10-13) but show different methylation patterns than the Denisovan (Extended Data Table 1, Fig. 1).

Note that we do not generally expect the number of DMRs along a lineage to be proportional to the length of the lineage, as this number is determined by several factors. First, the statistical power to detect DMRs depends on coverage and deamination levels. Thus, our ability to detect DMRs was lowest in the Denisovan, and highest in Ust’-Ishim. Second, the ability to filter out within-population variability was substantially higher along the AMH lineage, to which most samples belong. While filtering out such variability, we also exclude variability that exists across both AMH and archaic populations. This filtering also discards genomic regions that are variable between sexes, bone types and regions where methylation patterns tend to be more stochastic.

Variability that exists exclusively along the Neanderthal lineage was partially removed using the Vindija Neanderthal sample, which comes from a different bone (femur vs. phalanx) and age (adult vs. juvenile/adult). Along the Denisovan-lineage, on the other hand, such variability could not be filtered out using our array of samples (Fig. 1).

We also repeated the Gene ORGANizer analyses (see “Gene ORGANizer analysis” section) after removal of 20 DMRs that overlap regions which were shown to change methylation during osteogenic differentiation^55^. We show that the enrichment of voice-affecting genes holds, and thus, the differentiation state of cells in the samples is unlikely to explain the results we report.

### Comparison to previous reports

We have previously reported that compared to present-day humans, the HOXD cluster of genes is significantly hypermethylated in the Neanderthal and Denisovan samples^4^. Using the new methylation maps, we show that this observation holds (Extended Data Fig. 4b). Adding chimpanzee data, we see that similarly to AMHs, chimpanzee samples are also hypomethylated compared to archaic humans. This suggests that the hypermethylation arose along the archaic-human lineage. However, we find that the Ust’-Ishim individual is an outlier among modern humans, and that his methylation levels are closer to the Neanderthal than to modern humans, as was also shown by Hanghøj et al.^73^. The Neanderthal and Ust’-Ishim individuals are found >2 standard deviations from the mean observed methylation in modern humans. This suggests that although the Neanderthal is hypermethylated compared to almost all modern humans, she is not found completely outside modern human variation. The Denisovan, on the other hand, is found even further away, and significantly outside the other populations. Given this, the HOXD DMR was classified as Denisovan-derived (Extended Data Table 2). The Ust’-Ishim remains include a single femur, and to our knowledge, it was not compared morphologically to other humans.

Thus, further analysis is needed in order to determine whether the hypermethylation of the Ust’-Ishim individual compared to other AMHs is manifested in morphological changes as well.

Moreover, as this DMR is classified as Denisovan-derived, we cannot rule out the possibility that it is driven to some extent by age or bone type differences.

Compared to the previously reported DMRs^4^, in this study we found four times as many AMH- and archaic-derived DMRs (2,805 full bone DMRs compared to 891) and roughly twice as many Neanderthal- and Denisovan-derived DMRs (440 and 598 compared to 295 and 307 in the Denisovan and Neanderthal, respectively). The list of DMRs reported here cannot be directly compared to our previous list of DMRs because of several key differences in the analysis: (i) The previous study focused on DMRs that are invariable across tissues, whereas here we focused on DMRs in skeletal tissues. In the previous study, we were therefore able to extrapolate and find trends that extend beyond the skeletal system, such as neurological diseases. In this paper we focus on the skeletal system, hence the different appearance of the body map (Fig. 2b,c). (ii) the current study used stricter thresholds for DMR detection, including a minimum of 50 CpGs in each DMR (compared to 10 CpGs previously), and a requirement for physical overlap in the three-way DMR detection procedure. (iii) In this study, the AMH reference is a reconstructed ancient map, whereas in the previous study the AMH reference, as well as the other tissues used for filtering out noise, were mainly cultured cell lines with RRBS methylation maps.

When filtering DMRs along the lines of the previous study by taking only DMRs with low inter-tissue variability in humans (STD < 10%), we indeed observe similar trends. For example, when taking AMH-derived DMGs and analyzing their expression patterns using DAVID’s tissue expression tools^74^, we found that the brain is the most represented organ, with 51.5% of DMGs expressed in this organ (x1.28, FDR = 2.6 x 10^-4^), and glial cells are the most over-represented cell type (x20.6, but FDR > 0.05, UP_TISSUE DB, Extended Data Table 3). In fact, the brain is the only significantly enriched organ in this analysis. Similarly, when analyzing the GNF DB, we found that the subthalamic nucleus is the most enriched body part (x1.60, FDR = 9.2 x 10^-4^), followed by additional brain regions, such as the olfactory bulb (x1.54, FDR = 0.01), globus pallidus (x1.41, FDR = 0.04), and more (Extended Data Table 3). Similar enrichment patterns of the brain can be observed when analyzing expression patterns of all AMH-derived DMGs (Extended Data Table 3). Finally, we also find that similarly to the previous report, these DMRs are linked to diseases more often (23.1% compared to the genome average of 10.8%, DAVID OMIM_DISEASE DB^74^).

### Validation of face and larynx enrichment in Gene ORGANizer

To test whether the enrichment of the face and larynx could be attributed to the fact that the analyses are based on skeletal tissues, we tested whether the proportion of genes related to the face, larynx, vocal folds and pelvis within AMH-derived skeleton-related DMGs is higher than expected by chance. Out of 100 DMRs in genes that affect the skeleton, 31 genes are known to affect the voice, 34 affect the larynx, 87 affect the face, and 65 affect the pelvis, whereas genome-wide these proportions are significantly lower (14.2%, 20.2%, 70.0%, 52.4%, *P* = 1.0 x 10^-5^, *P* = 1.3 x 10^-3^, *P* = 2.1 x 10^-3^, P = 0.03, for vocal folds, larynx, face, and pelvis, respectively, hypergeometric test). For additional validation tests, see main text.

Genes associated with craniofacial features were taken from the GWAS-catalog (version 2019-04-21), using a threshold of *P* < 10^-8^. The following features were used: Dental caries, Cleft palate, Facial morphology, Intracranial volume, Cleft palate (environmental tobacco smoke interaction), Cranial base width, Craniofacial macrosomia, Facial morphology (factor 1, breadth of lateral portion of upper face), Facial morphology (factor 10, width of nasal floor), Facial morphology (factor 11, projection of the nose), Facial morphology (factor 12, vertical position of sublabial sulcus relative to central midface), Facial morphology (factor 14, intercanthal width), Lower facial height, Nose morphology, Nose size, Tooth agenesis (maxillary third molar), Tooth agenesis (third molar), facial morphology traits (multivariate analysis), Lower facial morphology traits (ordinal measurement), Lower facial morphology traits (quantitative measurement), Middle facial morphology traits (quantitative measurement), and Upper facial morphology traits (ordinal measurement). We then tested their overlap with DMGs. Genes associated with craniofacial features in the GWAS catalog significantly overlapped DMGs compared to the fraction expected by chance (5.17x, *P* = 3.4 x 10^-4^, hypergeometric test). As a control, we then tested how this 5-fold enrichment compares to non-craniofacial features. We used blood-related GWAS as a representative of general non-craniofacial GWAS. We extracted from the GWAS catalog 22 blood-related traits (the same number as extracted for craniofacial features), by taking the first 22 traits that appear in a search for the term “blood” and applying a threshold of *P* < 10^-8^. We then used these genes as a background control for the craniofacial enrichment. We observed a 3.86x enrichment of DMGs with regard to craniofacial-vs. non-craniofacial-associated genes (*P* = 0.011, chi-square test).

Additionally, we conducted a permutation test on the list of 129 AMH-derived DMGs that are linked to organs on Gene ORGANizer, replacing those that are linked to the skeleton with randomly selected skeleton-related genes. We then ran the list in Gene ORGANizer and computed the enrichment. We repeated the process 100,000 times and found that the enrichment levels we observed within AMH-derived DMGs are significantly higher than expected by chance for the laryngeal and facial regions, but not for the pelvis (*P* = 8.0 x 10^-5^, *P* = 3.6 x 10^-3^, *P* = 8.2 x 10^-4^, and *P* = 0.115, for vocal folds, larynx, face and pelvis, respectively, Extended Data Fig. 2b-e).

Potentially, longer genes have higher probability to overlap DMRs. Indeed, DMGs tend to be longer (148 kb vs. 39 kb, *P* = 9.9 x 10^-145^, *t*-test). We thus checked the possibility that genes affecting the larynx and face tend to be longer than other genes, and are thus more likely to contain DMRs. We found that length of genes could not be a factor explaining the enrichment within genes affecting the larynx, as these genes tend to be shorter than other genes in the genome (mean: 62.5 kb vs. 73.2 kb, *P* = 0.001, *t*-test). Genes affecting the face, on the other hand, tend to be longer than other genes (mean: 77.1 kb vs. 65.6 kb, *P* = 4.6×10^-5^, *t*-test). To examine if this factor may underlie the enrichment we observe, we repeated the analysis using only DMRs that are found within promoter regions (5 kb upstream to 1 kb downstream of TSS), thus eliminating the gene length factor. We found that the genes where such DMRs occur are still significantly associated with the face (*P* = 0.036, Fisher’s exact test). We next repeated the promoter DMR analysis for all genes and compared the Gene ORGANizer enrichment levels in this analysis to the genome-wide analysis. We observed very similar levels of enrichment (2.02x, 1.67x, and 1.24x, for vocal folds, larynx, and face, respectively, albeit FDR values > 0.05 due to low statistical power). Importantly, AMH-derived DMGs also do not tend to be longer than DMGs on the other branches (148 kb vs. 147 kb, *P* = 0.93, *t*-test). Together, these analyses suggest that gene length does not affect the observed enrichment in genes affecting the face and larynx.

Additionally, to test whether cellular composition or differentiation state could bias the results, we ran Gene ORGANizer on the list of DMGs, following the removal of 20 DMRs that are found <10 kb from loci where methylation was shown to change during osteogenic differentiation^55^. We found that genes affecting the voice and face are still the most over-represented (2.13x, 1.71x, and 1.27x, FDR = 0.032, FDR = 0.049, and FDR = 0.040, for vocal folds, larynx, and face, respectively, Extended Data Table 4).

We also investigated the possibility that (for an unknown reason) the DMR-detection algorithm introduces positional biases that preferentially identify DMRs within genes affecting the voice or face. To this end, we simulated stochastic deamination processes along the Ust’-Ishim, Altai Neanderthal, and Denisovan genomes, reconstructed methylation maps, and ran the DMR-detection algorithm on these maps. We repeated this process 100 times for each hominin and found no enrichment of any body part, including the face, vocal folds, or larynx (1.07x, 1.07x, and 1.04x, respectively, FDR = 0.88 for vocal folds, larynx, and face). Perhaps most importantly, none of the other archaic branches shows enrichment of the larynx or the vocal folds. However, archaic-derived DMGs show over-representation of the jaws, as well as the lips, limbs, scapulae, and spinal column (Extended Data Fig. 2f, Extended Data Table 4). In addition, DMRs that separate chimpanzees from all humans (archaic and modern, Extended Data Table 2) do not show over-representation of genes that affect the voice, larynx, or face, compatible with the notion that this trend emerged along the AMH lineage. We also sought to test whether the larynx and vocal folds, which we found to be significantly enriched only along the AMH lineage, are also enriched when compared to the other lineages. We ran a chi-squared test on the fraction of vocal folds- and larynx-affecting AMH-derived DMGs (25 and 29, respectively, out of a total of 120 organ-associated DMGs), compared to the corresponding fraction in the DMGs along all the other lineages (42 for vocal folds, 49 for larynx, out of a total of 275 organ-associated DMGs).

We found that both the larynx and vocal folds are significantly enriched in AMHs by over 50% compared to the other lineages (1.57x for both, P = 0.0248 and P = 0.0169 for vocal folds and larynx, respectively).

Furthermore, we added a human bone reduced representation bisulfite sequencing (RRBS) map, and produced a RRBS map from a chimpanzee infant unspecified long bone (Extended Data Table 1, see Methods). RRBS methylation maps include information on only ∼10% of CpG sites, and are biased towards unmethylated sites. Therefore, they were not included in the previous analyses. However, we added them in this part as they originate from a chimpanzee infant and a present-day human that is of similar age to the Denisovan (Extended Data Table 1), allowing sampling from individuals that are younger than the rest. Repeating the Gene ORGANizer analysis after including these samples in the filtering process, we found that the face and larynx are the only significantly enriched skeletal regions, and the enrichment within voice-affecting genes becomes even more pronounced (2.33x, FDR = 7.9 x 10^-3^, Extended Data Table 4).

We also examined if pleiotropy could underlie the observed enrichments. To a large extent, the statistical tests behind Gene ORGANizer inherently account for pleiotropy^11^, hence the conclusion that the most significant shared effect of the AMH-derived DMGs is in shaping vocal and facial anatomy is valid regardless of pleiotropy. Nevertheless, we tested this possibility more directly, estimating the pleiotropy of each gene by counting the number of different Human Phenotype Ontology (HPO) terms that are associated with it across the entire body^19^. We found that DMGs do not tend to be more pleiotropic than the rest of the genome (*P* = 0.17, *t*-test), nor do differentially methylated voice- and face-affecting genes tend to be more pleiotropic than other DMGs (*P* = 0.19 and *P* = 0.27, respectively).

Next, we tested whether the process of within-lineage removal of variable DMRs and the differential number of samples along each lineage biases the Gene ORGANizer enrichment analysis. To do so, we analyzed the pre-filtering DMRs along each lineage. We detect very similar trends to the post-filtering analysis, with the laryngeal and facial regions being the most significantly enriched within AMH-derived DMRs (1.58x, 1.44x and 1.21x-1.31x for the vocal folds, larynx and different facial regions, respectively, FDR < 0.05), and for archaic-derived DMRs, we detect no enrichment of the laryngeal region (FDR = 0.16 and FDR = 0.43 for the vocal folds and larynx, respectively), and the most enriched regions are the face, limbs, and urethra. With the exception of the urethra, these results are very similar to the results reported for the filtered DMRs, suggesting that the process of within-lineage removal of variable DMRs and the differential number of samples along each lineage does not bias the enrichment results.

Overall, we observe that AMH-derived DMGs across all 60 AMH samples are found outside archaic human variability, regardless of bone type, disease state, age, or sex, and that chimpanzee methylation levels in these DMGs cluster closer to archaic humans than to AMHs, suggesting that these factors are unlikely to underlie the observed trends.

Finally, we tested whether the filtering process in itself might underlie the observed trends. To this end, we re-ran the entire pipeline on Neanderthal- and Denisovan-derived DMGs, while applying to them all the filters as if they were Ust’-Ishim DMGs. This resulted in substantially fewer loci (89 for the Neanderthal and 50 for the Denisovan), which limits statistical power, but can still be used to examine whether there are any trends of enrichment similar to those observed in AMHs. We found no evidence that the filtering process could drive the enrichment of the vocal or facial areas: within Neanderthal-derived loci, filtered as if they were Ust’-Ishim-derived, we found that the vocal folds were ranked only 18th, with a non-significant enrichment of 1.27x (FDR = 0.815, compared to an enrichment of 2.11x within AMH-derived DMGs). The larynx was ranked 76th and showed a non-significant depletion of 0.87x (FDR = 0.783), and the face was ranked 31st, with a non-significant enrichment of 1.09x (FDR = 0.815). Within Denisovan-derived loci, filtered as if they were Ust’-Ishim-derived, none of the loci were linked to the vocal folds nor to the larynx (FDR = 0.535 and FDR = 0.834, respectively), and the face was ranked 30th (1.29x, FDR = 0.535, Extended Data Table 4). This test suggests that the filtering process in itself is very unlikely to underlie the enrichment of the vocal and facial parts within AMH-derived DMGs.

Next, we applied the Neanderthal/Denisovan filters to the Ust’-Ishim-derived loci. This resulted in 792 loci. We found that the vocal folds remained the most enriched body part (1.76x, FDR = 0.032), the larynx was marginally significant (1.53x, FDR = 0.0502), and the facial region was significantly enriched too (e.g., cheek and chin ranked 2nd, 3rd within significantly enriched body parts, 1.66x and 1.63x, FDR = 0.031 and FDR = 0.013, respectively, Extended Data Table 4). Importantly, we do not rule out the option that extensive regulatory changes in genes related to vocal and facial anatomy might have occurred along the Neanderthal and Denisovan lineages as well. Indeed, as we report in Extended Data Fig. 2, parts of the face are enriched within Archaic-derived DMGs. However, we currently see no substantial evidence supporting this.

Importantly, the link between genetic alterations and phenotypes related to the voice is complex. Some brain-related disorders (i.e., clinical disorders that affect the brain) result in alterations to the voice, the mechanism in which is very difficult to pin down. Although the mechanism leading to voice alterations (either in its pitch, timbre, volume or range) in some of the genes we report is unknown, many of the disorders are skeletal, suggesting the mechanism is related to anatomical changes to the vocal tract. Such changes could also affect more primary functions of the larynx, such as swallowing and breathing. However, the enrichment we observe in Gene ORGANizer shows these genes were also shown to drive vocal alterations in the disorders they underlie^11, 19^. Voice and speech alterations were also shown to be driven by cultural, dietary and behavioral changes affecting bite configuration^75^. Here too, these factors are unlikely to underlie the vocal alterations in the genes we report, as individuals from the same family as the individual with the disorder, who do not carry the dysfunctional allele, were not reported to present any vocal phenotypes.

The larynx is an organ which is primarily involved in breathing and swallowing in mammals. In humans, the larynx is also used to produce complex speech, but not every change to the larynx necessarily affects speech. Despite these additional functions, the genes reported by Gene ORGANizer and HPO were specifically associated with voice alterations, directly or indirectly, suggesting that although they could have additional effects, their effect on the voice is their most shared function.

### Computing correlation between methylation and expression

In order to identify regions where DNA methylation is tightly linked with expression levels, we scanned each DMR in overlapping windows of 25 CpGs (the window used for smoothing the deamination signal). In each window we computed the correlation between DNA methylation levels and expression levels of overlapping genes as well as the closest genes upstream and downstream genes, across 21 tissues^32^. For each DMR, we picked the window with the best correlation (in absolute value) and computed regression FDR-adjusted *P*-value. DMRs that overlap windows with FDR < 0.05 were considered to be regions where methylation levels are significantly correlated with expression levels. 90 such DMRs were found among the skeletal AMH-derived DMRs, 93 among the archaic human-derived DMRs, 40 among Neanderthal-derived DMRs, and 19 among Denisovan-derived DMRs.

As no expression data were available for Ust’-Ishim, Bone1 and Bone2, we approximated their *NFIX* expression level by taking the average of *NFIX* expression from three osteoblast RNA-seq datasets that were downloaded from GEO accession numbers GSE55282, GSE85761 and GSE78608. RNA-seq data for chondrocytes was downloaded from the ENCODE project, GEO accession number GSE78607 and plotted against measured methylation levels in primary chondrocytes (see “Human primary chondrocyte validation” chapter). Notably, even though the expression and methylation data come from different individuals, plotting them against one another positions them only ∼one standard deviation from the expression value predicted by the regression line (Fig. 5b). Future studies providing RNA expression levels for the laryngeal skeleton and vocal folds might provide further information on the methylation-expression links of these genes.

### Studying the function of DMGs

#### Gene Ontology analysis

Gene ontology and expression analyses were conducted using Biological Process and UNIGENE expression tools in DAVID^74^, using an FDR threshold of 0.05.

#### Gene ORGANizer analysis

Similarly to sequence mutations, changes in regulation are likely to be unequally distributed across different body systems, owing to negative and positive selection, as well as inherent traits of the genes affecting each organ. Thus, we turned to investigate which body parts are affected by the DMGs. To this end, we ran the lists of DMGs in Gene ORGANizer^11^, which is a tool that links genes to the organs they affect, through known disease and normal phenotypes. Thus, it allows us to investigate directly the phenotypic function of genes, to identify their shared targets and to statistically test the significance of such enrichments. We ran the lists of DMGs in the ORGANize option using the default parameters (i.e., based on *confident* and *typical* gene-phenotype associations).

When we ran the list of skeletal AMH-derived DMRs, we found 11 significantly enriched body parts, with the vocal folds and the larynx being the most enriched parts (x2.11 and x1.68, FDR = 0.017 and FDR = 0.048, respectively). Most other parts belonged to the face (teeth, forehead, lips, eyelid, maxilla, face, jaws), as well as the pelvis and nails (Fig. 2c,d, Extended Data Table 4). For archaic-derived DMGs, the lips, limbs, jaws, scapula, and spinal column were enriched (Extended Data Fig. 2f, Extended Data Table 4). The Neanderthal-derived and Denisovan-derived DMG lists did not produce any significantly enriched organs, but the immune system was significantly depleted within Neanderthal-derived DMRs (x0.67, FDR = 0.040).

In order to examine whether such trends could arise randomly from the reconstruction method, we repeated the analysis on the previously described 100 simulations. We ran all simulated DMGs (4,153) in Gene ORGANizer and found that no enrichment was detected, neither for voice-related organs (vocal folds: x0.99, FDR = 0.731, larynx: x.1.02, FDR = 0.966, FDR = 0.966), nor for any other organ.

#### Overlap with enhancer regions

To further test whether the AMH-derived DMRs overlap skeletal regulatory regions, we examined the previously reported 403,968 human loci, where an enrichment of the active enhancer mark H3K27ac was detected in developing human limbs (E33, E41, E44, and E47)^76^. Each DMR was allocated a random genomic position in its original chromosome, while keeping its original length and matching the distribution of GC-content and CpG density between the original and permutated lists. GC-content and CpG density matching was done by matching a 10-bin histogram of the original and permutated lists. This was repeated for 10,000 iterations.

We found that AMH-derived DMRs overlap limb H3K27ac-enriched regions ∼2x more often than expected by chance (610 overlapping DMRs, compared to 312.4±21.7, *P* < 10^-4^, permutation test).

SOX9 upstream putative enhancer coordinates used in Fig. 4b were taken from ^13, 14, 23, 77, 78^.

### Computing the density of changes along the genome

We computed the density of derived CpG positions along the genome in two ways. First, we used a 100 kb window centered in the middle of each DMR and computed the fraction of CpGs in that window which are differentially methylated (i.e., are found within a DMR). Second, for the chromosome density plots, we did not center the window around each DMR, but rather used a non-overlapping sliding 100 kb window starting at position 1 and running the length of the chromosome.

### NFIX, COL2A1, SOX9, ACAN and XYLT1 phenotypes

The vocal tract and larynx affecting genes presented in this paper show involvement in laryngeal cartilage and soft tissue phenotypic variation. Clinical phenotypes can be of high severity, with substantial impacts on normal breathing functions, to the point where the cause of death is due to respiratory distress. *SOX9* and *NFIX* are often associated with laryngomalacia^11, 19^ (Extended Data Table 5), a collapse of the larynx due to malformation of the laryngeal cartilaginous framework and/or malformed connective tissues, particularly during inhalation. Patients with mutations in *COL2A1* often show backwards displacement of the tongue base^11, 19^. Less severe phenotypes of the reported genes include variation of voice quality in the form of pitch variation (high in patients suffering from *XYLT1* mutations) and sometimes hoarseness of the voice (reported for some patients with mutations of *ACAN*, Extended Data Table 5)^11, 19^. Whether this is due to variation of the vocal tract and laryngeal anatomy influenced by the *ACAN* mutation or due to a scaled down vocal tract size in the case of the *XYLT1* mutation which also causes primordial dwarfism is not yet clear.

#### NFIX phenotypes

Skeletal phenotypes that are associated with the Marshall-Smith syndrome were extracted from the Human Phenotype Ontology (HPO) ^19^. Non-directional phenotypes (e.g., irregular dentition) and phenotypes that are expressed in both directions (e.g., tall stature and short stature) were removed.

Mutations in *NFIX* have also been linked to the Sotos syndrome. However, *NFIX* is not the only gene that was linked to this syndrome; mutations in *NSD1* were also shown to drive similar phenotypes^51^. Therefore, it is less relevant in assessing the functional consequences of general shifts in the activity levels of *NFIX*. Nevertheless, it is noteworthy that in the Sotos syndrome too, most symptoms are a mirror image of the Neanderthal phenotype (e.g., prominent chin and high forehead).

Comparing of SOX9, ACAN, COL2A1, and NFIX expression between AMH and mouse 93 appendicular skeleton samples were used to compare expression levels of *NFIX*, *SOX9*, *ACAN* and *COL2A1* in human and mouse: 1. Five Human expression array data of iliac bones ^79^, downloaded from ArrayExpress accession number E-MEXP-2219. 2. 84 Human expression array of iliac bones, downloaded from ArrayExpress accession number E-MEXP-1618. 3. Three Mouse expression array data of femur and tibia bones, downloaded from ArrayExpress accession number E-GEOD-61146. 4. One Mouse RNA-seq of a tibia bone, downloaded from supplementary data. Expression values were converted to percentiles, according to each gene expression level compared to the rest of the genome across each sample (Fig. 5c).

### Comparing *SOX9, ACAN, COL2A1, XYLT1*, and *NFIX* methylation between AMH, chimpanzee and Neanderthal femora

To check whether the AMH hypermethylation of *SOX9, ACAN, COL2A1, XYLT1* and *NFIX* could be a result of variability between bone types, we compared the four chimpanzee femur 850K methylation arrays to the 52 present-day femur 450K methylation arrays. We took probes within AMH-derived DMRs that appear on both arrays. We found that these genes are consistently hypermethylated in AMHs (*P =* 1.6×10^-7^, t-test), with 38 probes showing >5% hypermethylation in AMH, whereas only eight probes show such hypermethylation in chimpanzees (Extended Data Fig. 3d). Therefore, even when comparing methylation from the same bone, same sex, same developmental stage, measured by the same technology, and across the same positions, AMH show consistent hypermethylation across all of these DMGs.

Similarly, when comparing the DMRs in *SOX9*, *ACAN*, *COL2A1*, *XYLT1*, and *NFIX* between the Ust’-Ishim and Vindija Neanderthal samples, the Vindija Neanderthal sample is consistently hypomethylated compared to the Ust’-Ishim individual (*P* = 1.2 x 10^-5^, Extended Data Fig. 3c). Both of these samples were extracted from femora of adult individuals, and methylation was reconstructed using the same technology. This suggests that the hypermethylation of AMHs compared to Neanderthals is unlikely to be driven by age or bone type, and rather reflects evolutionary shifts.

### Scanning the SOX9 region for mutations altering NFI binding motifs

To examine whether the changes in regulation of *SOX9* could possibly be explained by changes in the binding sites of NFI proteins, we searched for the NFI motif ^80, 81^ along the gene body and the 350 kb upstream region of *SOX9*. We looked for NFI motifs that exist in the genomes of the Altai and Vindija Neanderthal, as well as in the Denisovan, but were abolished in AMHs. We did not find any evidence of such substitutions.

### Comparison to divergent traits between Neanderthals and AMHs

To further investigate potential phenotypic consequences of the DMGs we report, we probed the HPO database^19^ and compared these HPO phenotypes to known morphological differences between Neanderthals and modern humans. To compile a list of traits in which Neanderthals and AMHs differ, we reviewed key sources that surveyed Neanderthal morphology summarized in ^12^. We identified traits in which Neanderthals are found completely outside AMH variation, as well as traits where one group is significantly different from the other, but the distribution of observed measurements partially overlap. Non-directional traits (i.e., traits that could not be described on scales such as higher/lower, accelerated/delayed etc.) were not included, as could not be paralleled with HPO phenotypes. The compiled list included 107 phenotypes, 75 of which have at least one equivalent HPO phenotype (4.8 on average). For example, the HPO phenotype Taurodontia (HP:0000679) was linked to the trait “Taurodontia”, and the following HPO phenotypes were linked to the trait “Rounded and robust rib shafts”: *Broad ribs* (HP:0000885), *Hypoplasia of first ribs* (HP:0006657), *Short ribs* (HP:0000773), *Thickened cortex of long bones* (HP:0000935), *Thickened ribs* (HP:0000900), *Thin ribs* (HP:0000883), *Thoracic hypoplasia* (HP:0005257). For each skeleton-affecting phenotype, we determined whether it matches a known morphological difference between Neanderthals and AMHs. For example, *Hypoplastic ilia* (HPO ID: HP:0000946) was marked as divergent because in the Neanderthal the iliac bones are considerably enlarged compared to AMHs^12^. We then counted for each gene (whether DMG or not) the fraction of its associated HPO phenotypes that are divergent between Neanderthals and AMHs. We found that four of the top five most differentially methylated skeletal genes (*XYLT1*, *NFIX*, *ACAN*, and *COL2A1*) are in the top 100 genes with the highest fraction of divergent traits between Neanderthals and AMHs (out of a total of 1,789 skeleton-related genes). In fact, *COL2A1*, which is the top ranked DMR (Extended Data Table 2), is also the gene that is overall associated with the highest number of derived traits (63) (Extended Data Table 7). This suggests that these extensive methylation changes are possibly linked to phenotypic divergence between archaic and AMHs.

## Data and Software Availability

All methylation data generated in this work have been deposited in NCBI’s Gene Expression Omnibus under GEO accession number <u>GSE96833</u>.

## Extended Data Figures

**Extended Data Figure 1.**
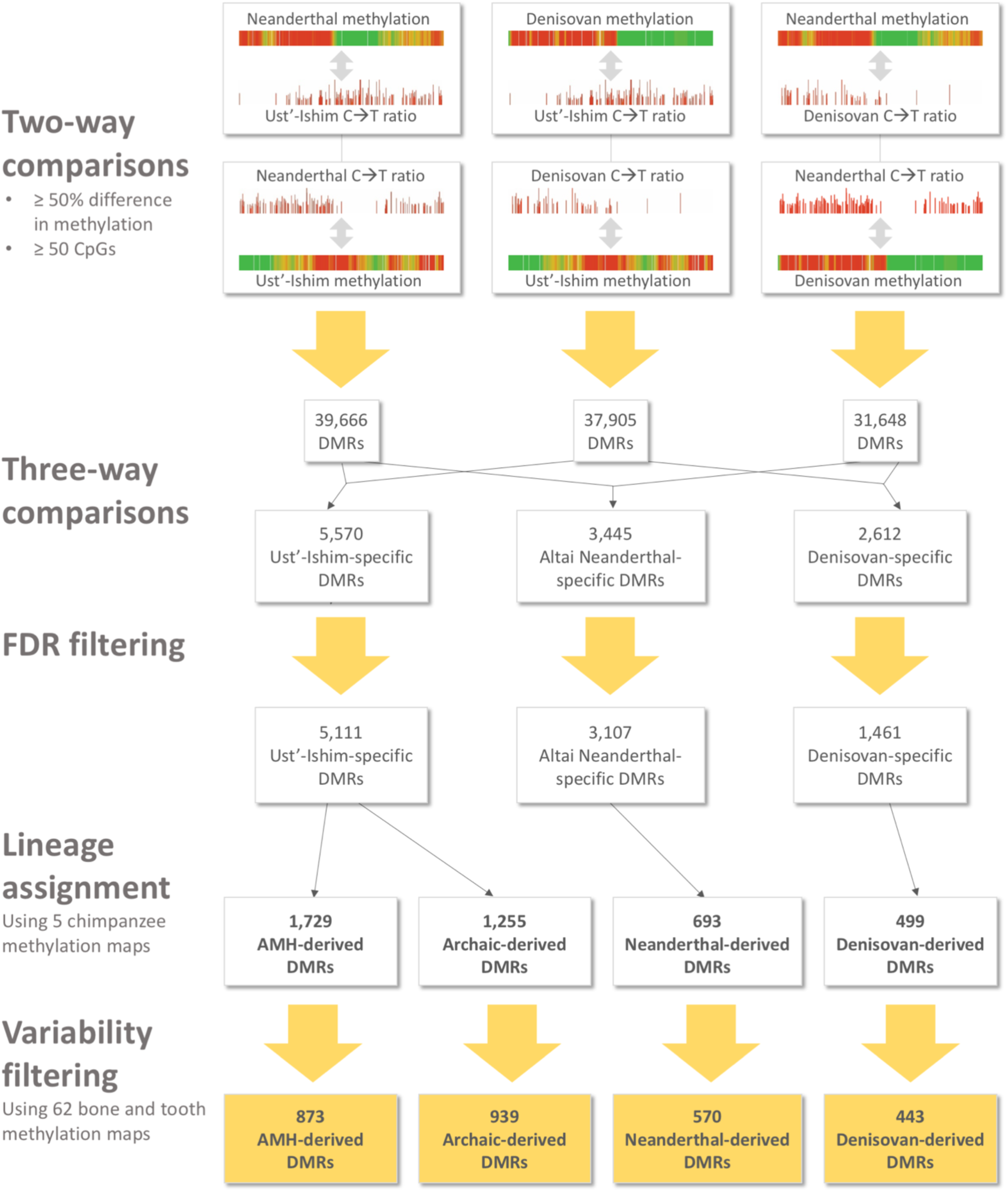
DMR-detection flowchart. At the core of the process are six two-way (pairwise) comparisons between the Altai Neanderthal, Denisovan, and Ust’-Ishim individuals. In each two-way comparison, a C→T deamination signal of one hominin was compared to the reconstructed methylation map of the other hominin. This resulted in three lists of pairwise DMRs, that were then intersected to identify hominin-specific DMRs, defined as DMRs that appear in two of the lists. False discovery rates were controlled by running 100 simulations for each hominin, each simulating the processes of deamination, methylation reconstruction, and DMR-detection. Only DMRs that passed FDR thresholds of < 0.05 were kept (see Methods). To discard non-evolutionary DMRs we used 62 skeletal methylation maps, and kept only loci whose methylation levels differed in one lineage, regardless of age, bone type, disease or sex. Finally, five chimpanzee methylation maps were used to assign the lineage in which each DMR likely emerged.

**Extended Data Figure 2.**
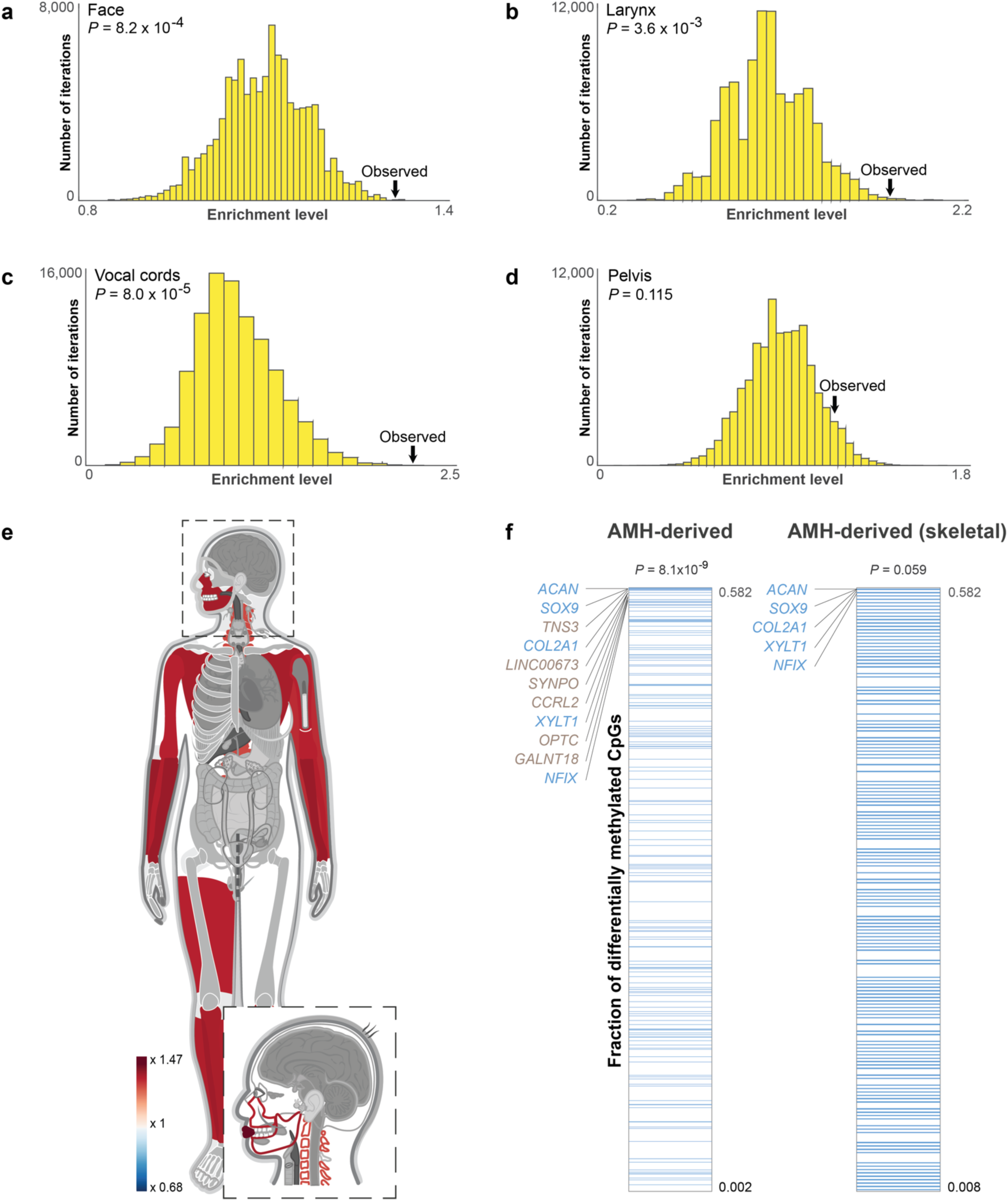
The face and larynx are enriched within AMH-derived DMGs compared to genes affecting the skeleton, and compared to archaic-derived DMGs. **a-d.** The distribution of enrichment levels in 100,000 randomized lists of genes, where non-skeletal AMH-derived DMGs were unchanged, whereas skeleton-related DMGs were replaced with random skeleton-related genes. Observed enrichment levels are significantly higher than expected in the face, larynx, and vocal folds. **e.** A heat map representing the level of enrichment of each anatomical part within archaic-derived DMGs. Genes affecting the lips, limbs, jaws, scapula, and spinal column are the most enriched within archaic-derived DMRs. Only body parts that are significantly enriched (FDR < 0.05) are colored. **f.** The number of AMH-derived CpGs per 100 kb centered around the middle of each DMR. Genes were ranked according to the fraction of derived CpG positions within them. Genes affecting the face are marked with blue lines. AMH-derived DMGs which affect the face tend to be ranked significantly higher. Although only ∼2% of genes in the genome are known to affect lower and midfacial projection, three of the top five AMH-derived DMGs, and all top five AMH-derived skeleton-affecting DMGs affect facial projection.

**Extended Data Figure 3.**
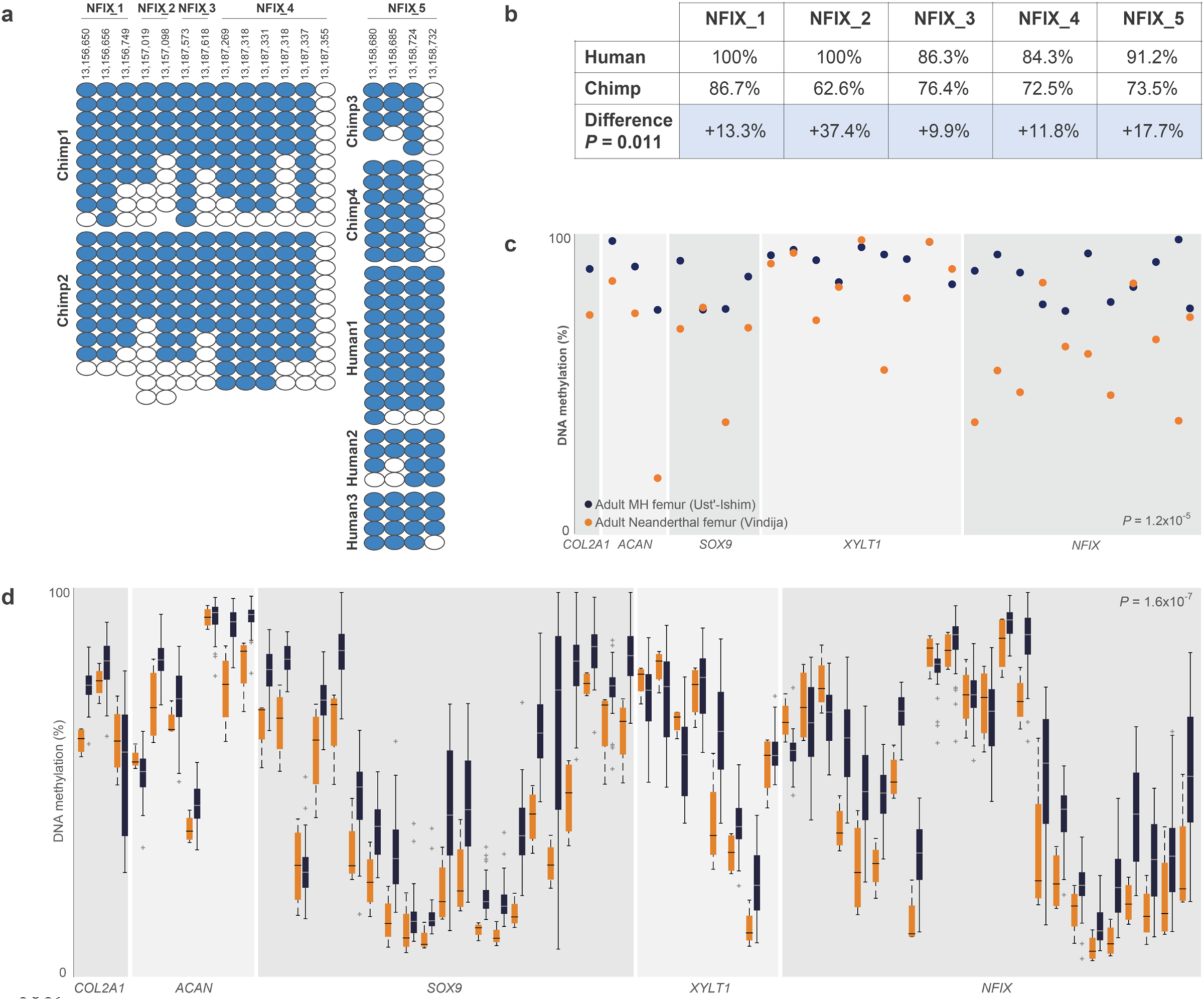
AMHs are hypermethylated compared to Neanderthal and chimpanzee bone samples, even when age and bone type are considered. **a,b.** Bisulfite-PCR in human and chimpanzee crania of five regions within the two *NFIX* DMRs, showing hypermethylation of NFIX in AMHs (*P* = 0.011, *t*-test). Each column represents a CpG position, with each circle representing either methylated (blue) or unmethylated (white) measurements. Human hg19 coordinates are shown for each CpG position. Chimpanzee methylation in regions 1-4 was compared with the human I1583 cranium. Region 5 was compared with I1583 and three additional present-day human crania, presented in the figure. Summarized results are presented in the table. **c.** *COL2A1, ACAN, SOX9,* and *NFIX* are hypermethylated in Ust’-Ishim (blue) compared to the Vindija Neanderthal (orange). Circles represent mean methylation levels in AMH-derived DMRs. Both samples were extracted from femora of adults, and methylation was reconstructed using the same method. The DMRs presented include also those that were analyzed in the density analyses (see Methods). The hypermethylation of these genes in AMHs is unlikely to be attributed to age or bone type. **d.** *COL2A1, ACAN, SOX9,* and *NFIX* are hypermethylated in AMH femora compared to chimpanzee femora. Each pair of box plots represents methylation levels across 52 AMH femora (blue) and four chimpanzee femora (orange) in a single probe of methylation array. When comparing methylation in the same bone, measured by the same technology, and across the same positions, AMHs show almost consistent hypermethylation compared to chimpanzee. The probes presented include also probes within DMRs that were analyzed in the density analyses (see Methods).

**Extended Data Figure 4.**
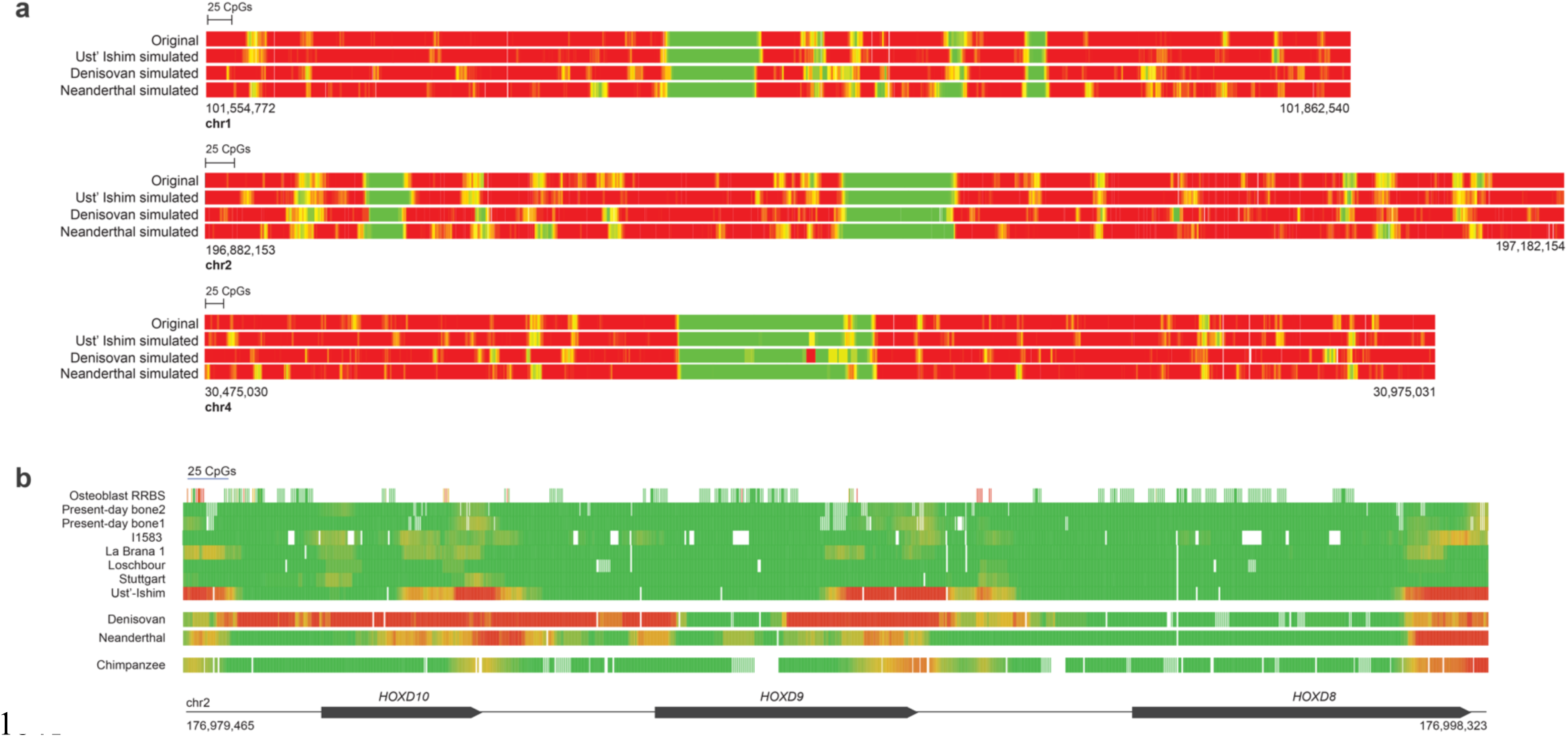
Simulations of deamination and reconstruction, and comparison to previous reports. **a.** Simulations of cytosine deamination, followed by reconstruction reproduce DNA methylation maps. Deamination was simulated for each position based on its methylation level, read coverage and the observed rate of deamination in each hominin. Then, DNA methylation maps were reconstructed and matched against the original map. The number of DMRs found were used as an estimate of false discovery rate. Three exemplary regions are presented, where methylation levels are color-coded from green (unmethylated) to red (methylated). **b.** The HOXD cluster is hypermethylated in archaic humans, and in the Ust’-Ishim individual. Methylation levels are color-coded from green (unmethylated) to red (methylated). The top eight bars show ancient and present-day AMH samples, the lower three show the Denisovan, Neanderthal and chimpanzee. The promoter region of *HOXD9* is hypermethylated in the Neanderthal and the Denisovan, but not in AMHs. The 3’ ends of the three genes are hypermethylated in the Neanderthal, Denisovan, Ust’-Ishim and chimpanzee, but not in other AMH samples. The promoter of HOXD10 is methylated only in the Denisovan.

**Extended Data Figure 5.**
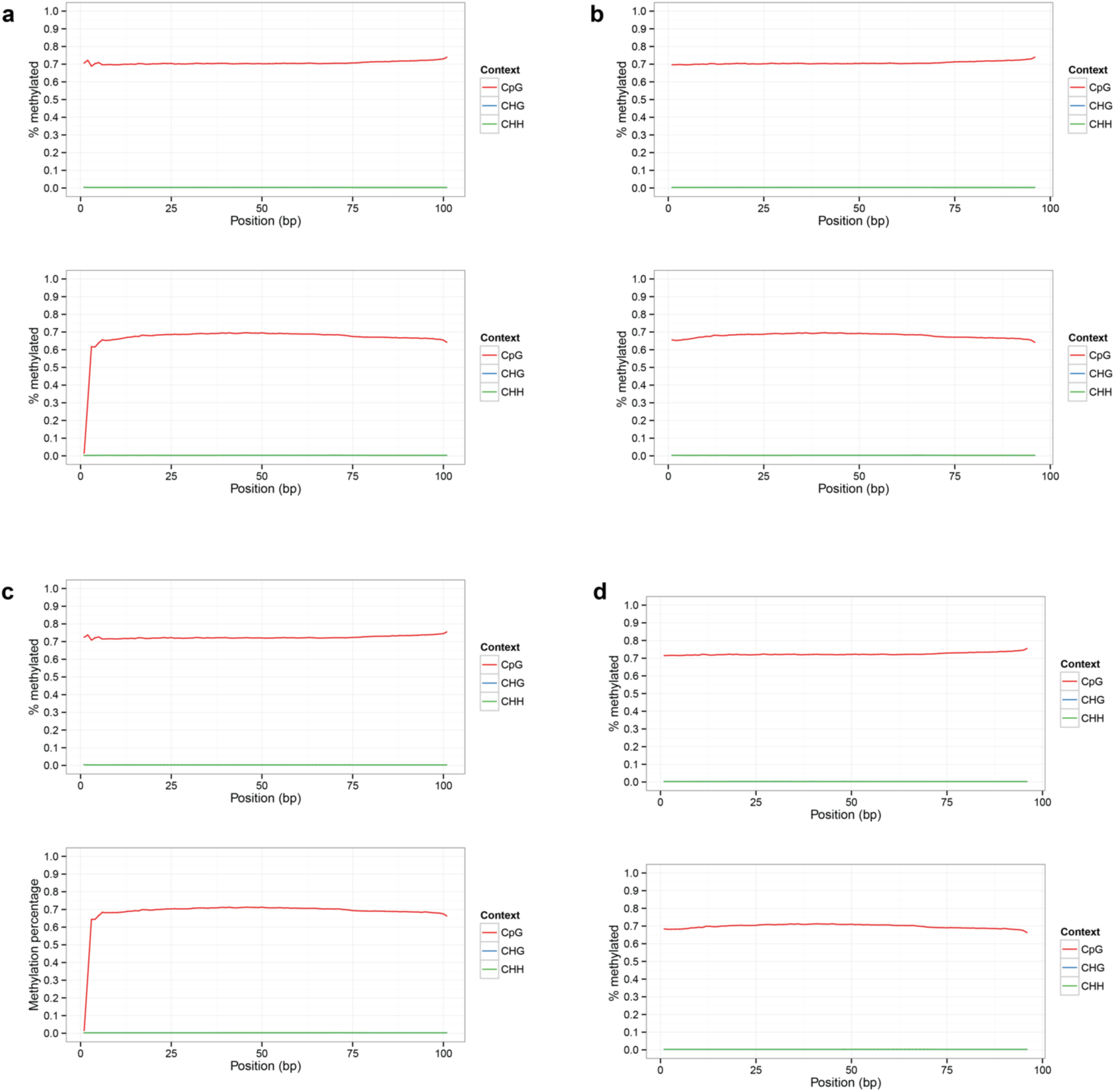
M-bias plots along reads in bone sample 1 and sample 2. **a**. Pre-filtering methylation along read1 and read2 in the autosomes of bone 1. **b**. Post-filtering methylation along read1 and read2 in the autosomes of bone 1. **c**. Pre-filtering methylation along read1 and read2 in the autosomes of bone 2. **d**. Post-filtering methylation along read1 and read2 in the autosomes of bone 2.

**Extended Data Figure 6.**
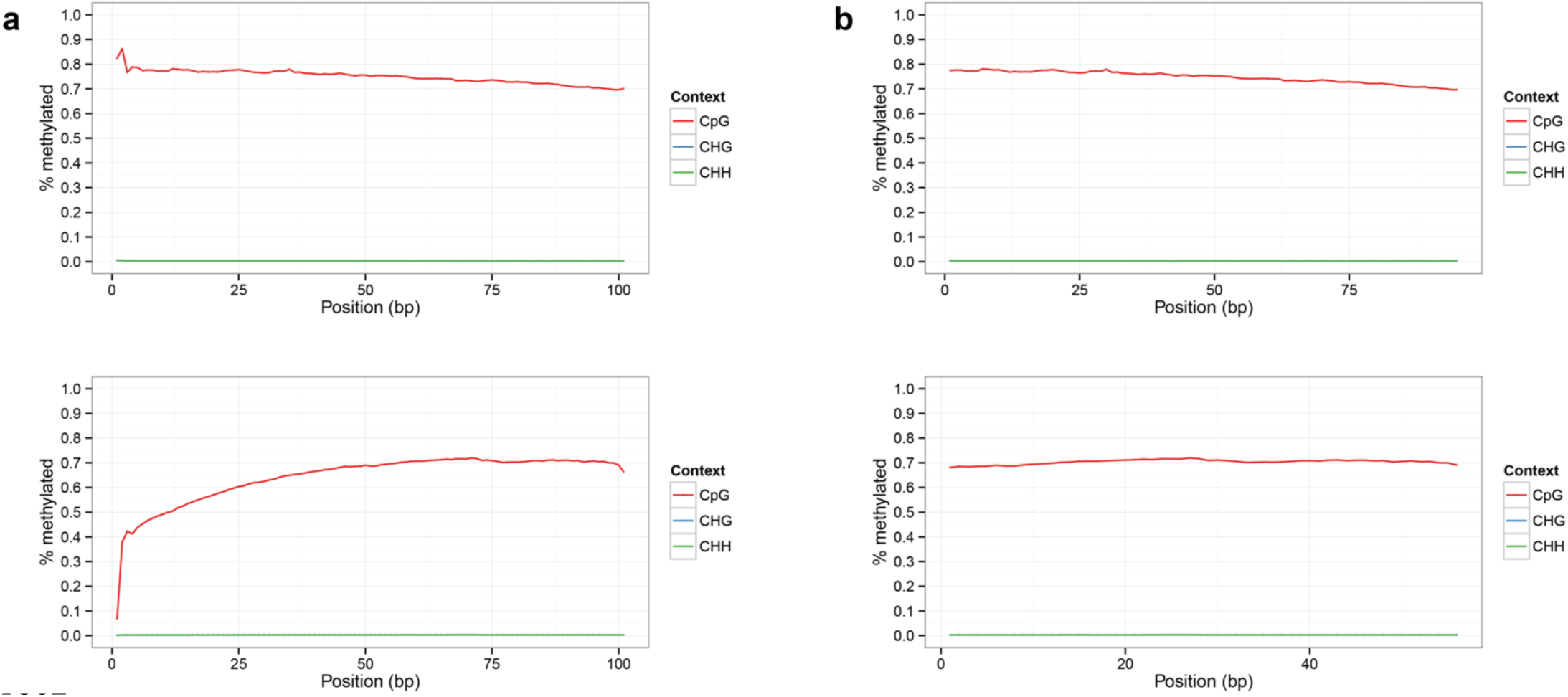
M-bias plots along reads in the chimpanzee rib sample. **a**. Pre-filtering methylation along read1 and read2 in the autosomes. **b**. Post-filtering methylation along read1 and read2 in the autosomes.

## References

1. Prüfer, K. et al. The complete genome sequence of a Neanderthal from the Altai Mountains. Nature 505, 43–9 (2014).

2. King, M. C. & Wilson, A. C. Evolution at two levels in humans and chimpanzees. Science 188, 107–116 (1975).

3. Hernando-Herraez, I., Garcia-Perez, R., Sharp, A. J. & Marques-Bonet, T. DNA Methylation: Insights into Human Evolution. PLoS Genetics 11, e1005661 (2015).

4. Gokhman, D. et al. Reconstructing the DNA methylation maps of the Neandertal and the Denisovan. Science 344, 523–527 (2014).

5. Prüfer, K. et al. A high-coverage Neandertal genome from Vindija Cave in Croatia. Science (80-.). 358, 655–658 (2017).

6. Fu, Q. et al. Genome sequence of a 45,000-year-old modern human from western Siberia. Nature 514, 445–9 (2014).

7. Lazaridis, I. et al. Ancient human genomes suggest three ancestral populations for present-day Europeans. Nature 513, 409–413 (2014).

8. Olalde, I. et al. Derived immune and ancestral pigmentation alleles in a 7,000-year-old Mesolithic European. Nature 507, 225–228 (2014).

9. Mathieson, I. et al. Genome-wide patterns of selection in 230 ancient Eurasians. Nature 528, 499– 503 (2015).

10. Gokhman, D., Malul, A. & Carmel, L. Inferring Past Environments from Ancient Epigenomes. Mol. Biol. Evol. 34, 2429–2438 (2017).

11. Gokhman, D. et al. Gene ORGANizer: Linking genes to the organs they affect. Nucleic Acids Res. 45, W138–W145 (2017).

12. Aiello, L. & Dean, C. An Introduction to Human Evolutionary Anatomy. (Elsevier, 2002).

13. Prescott, S. L. et al. Enhancer Divergence and cis-Regulatory Evolution in the Human and Chimp Neural Crest. Cell 163, 68–84 (2015).

14. Wilderman, A., VanOudenhove, J., Kron, J., Noonan, J. P. & Cotney, J. High-Resolution Epigenomic Atlas of Human Embryonic Craniofacial Development. Cell Rep. 23, 1581–1597 (2018).

15. MacArthur, J. et al. The new NHGRI-EBI Catalog of published genome-wide association studies (GWAS Catalog). Nucleic Acids Res. 45, D896–D901 (2017).

16. Hernando-Herraez, I. et al. The interplay between DNA methylation and sequence divergence in recent human evolution. Nucleic Acids Res 43, 8204–8214 (2015).

17. Schultz, M. D. et al. Human body epigenome maps reveal noncanonical DNA methylation variation. Nature 523, 212–6 (2015).

18. Peyrégne, S., Boyle, M. J., Dannemann, M. & Prüfer, K. Detecting ancient positive selection in humans using extended lineage sorting. Genome Res. 27, 1563–1572 (2017).

19. Köhler, S. et al. The Human Phenotype Ontology project: Linking molecular biology and disease through phenotype data. Nucleic Acids Res. 42, (2014).

20. Lee, Y. H. & Saint-Jeannet, J. P. Sox9 function in craniofacial development and disease. Genesis 49, 200–208 (2011).

21. Ng, L. J. et al. Sox9 binds DNA, activates transcription, and coexpresses with type II collagen during chondrogenesis in the mouse. Dev. Biol. 183, 108–121 (1997).

22. Claes, P. et al. Genome-wide mapping of global-to-local genetic effects on human facial shape. Nat. Genet. 50, 414–423 (2018).

23. Yao, B. et al. The SOX9 upstream region prone to chromosomal aberrations causing campomelic dysplasia contains multiple cartilage enhancers. Nucleic Acids Res. 43, 5394–5408 (2015).

24. Kim, K. Il, Park, Y. S. & Im, G. Il. Changes in the epigenetic status of the SOX-9 promoter in human osteoarthritic cartilage. J. Bone Miner. Res. 28, 1050–1060 (2013).

25. Claes, P. et al. Modeling 3D Facial Shape from DNA. PLoS Genet. 10, e1004224 (2014).

26. Cheng, P. F. et al. Methylation-dependent SOX9 expression mediates invasion in human melanoma cells and is a negative prognostic factor in advanced melanoma. Genome Biol. 16, 42 (2015).

27. Huang, C.-Z. et al. Sox9 transcriptionally regulates Wnt signaling in intestinal epithelial stem cells in hypomethylated crypts in the diabetic state. Stem Cell Res. Ther. 8, 60 (2017).

28. Pamnani, M., Sinha, P., Singh, A., Nara, S. & Sachan, M. Methylation of the Sox9 and Oct4 promoters and its correlation with gene expression during testicular development in the laboratory mouse. Genet. Mol. Biol. 39, 452–458 (2016).

29. Hall, B. D. Lethality in Desbuquois dysplasia: Three new cases. Pediatr. Radiol. 31, 43–47 (2001).

30. LaCroix, A. J. et al. GGC Repeat Expansion and Exon 1 Methylation of XYLT1 Is a Common Pathogenic Variant in Baratela-Scott Syndrome. Am. J. Hum. Genet. (2018). doi:https://doi.org/10.1016/j.ajhg.2018.11.005

31. Ohba, S., He, X., Hojo, H. & McMahon, A. P. Distinct Transcriptional Programs Underlie Sox9 Regulation of the Mammalian Chondrocyte. Cell Rep. 12, 229–243 (2015).

32. Roadmap Epigenomics Consortium et al. Integrative analysis of 111 reference human epigenomes. Nature 518, 317–329 (2015).

33. Carrió, E. et al. Deconstruction of DNA methylation patterns during myogenesis reveals specific epigenetic events in the establishment of the skeletal muscle lineage. Stem Cells 33, 2025–2036 (2015).

34. Maunakea, A. K. et al. Conserved role of intragenic DNA methylation in regulating alternative promoters. Nature 466, 253–7 (2010).

35. Ford, E. E. et al. Frequent lack of repressive capacity of promoter DNA methylation identified through genome-wide epigenomic manipulation. bioRxiv (2018). doi:https://doi.org/10.1101/170506

36. Housman, G., Havill, L. M., Quillen, E. E., Comuzzie, A. G. & Stone, A. C. Assessment of DNA Methylation Patterns in the Bone and Cartilage of a Nonhuman Primate Model of Osteoarthritis. Cartilage (2018). doi:10.1177/1947603518759173

37. Pjanic, M. et al. Nuclear Factor I genomic binding associates with chromatin boundaries. BMC Genomics 14, 99 (2013).

38. Tompson, S. W., et al. A Recessive Skeletal Dysplasia, SEMD Aggrecan Type, Results from a Missense Mutation Affecting the C-Type Lectin Domain of Aggrecan. Am. J. Hum. Genet. 84, 72– 79 (2009).

39. Frenzel, K., Amann, G. & Lubec, B. Deficiency of laryngeal collagen type II in an infant with respiratory problems. Arch. Dis. Child. 78, 557–9 (1998).

40. Hoornaert, K. P. et al. Stickler syndrome caused by COL2A1 mutations: genotype–phenotype correlation in a series of 100 patients. Eur J Hum Genet. 18, 872–880 (2010).

41. Shaw, A. C. et al. Phenotype and natural history in Marshall-Smith syndrome. Am. J. Med. Genet. Part A 152, 2714–2726 (2010).

42. Lieberman, P. The Evolution of Human Speech: Its Anatomical and Neural Bases. Curr. Anthropol. 48, 39–66 (2007).

43. Fitch, W. T., de Boer, B., Mathur, N. & A. Ghazanfar, A. Response to Lieberman on “Monkey vocal tracts are speech-ready”. Sci. Adv. 3, (2017).

44. Lieberman, P. Comment on “Monkey vocal tracts are speech-ready”. Sci. Adv. 3, (2017).

45. Nishimura, T. Developmental changes in the shape of the supralaryngeal vocal tract in Chimpanzees. Am. J. Phys. Anthropol. (2005). doi:10.1002/ajpa.20112

46. De Boer, B. Modelling vocal anatomy’s significant effect on speech. *J*. Evol. Psychol. 8, 351–366 (2010).

47. Lieberman, D. E. The Evolution of the Human Head. (Harvard University Press, 2011).

48. Nishimura, T., Mikami, A., Suzuki, J. & Matsuzawa, T. Descent of the hyoid in chimpanzees: evolution of face flattening and speech. J. Hum. Evol. 51, 244–254 (2006).

49. Fitch, W. T. The evolution of speech: A comparative review. Trends in Cognitive Sciences 4, 258–267 (2000).

50. Steele, J., Clegg, M. & Martelli, S. Comparative morphology of the hominin and african ape hyoid bone, a possible marker of the evolution of speech. Hum Biol 85, 639–672 (2013).

51. Malan, V. et al. Distinct effects of allelic NFIX mutations on nonsense-mediated mRNA decay engender either a sotos-like or a Marshall-Smith Syndrome. Am. J. Hum. Genet. 87, 189–198 (2010).

52. Van Balkom, I. D. C. et al. Development and behaviour in Marshall-Smith syndrome: An exploratory study of cognition, phenotype and autism. J. Intellect. Disabil. Res. (2011). doi:10.1111/j.1365-2788.2011.01451.x

53. Cullen, A., Clarke, T. A. & O’Dwyer, T. P. The Marshall-Smith syndrome: a review of the laryngeal complications. Eur J Pediatr 156, 463–464 (1997).

54. Ziller, M. J. et al. Charting a dynamic DNA methylation landscape of the human genome. Nature 500, 477–481 (2013).

55. Håkelien, A. M. et al. The regulatory landscape of osteogenic differentiation. Stem Cells 32, 2780–2793 (2014).

56. Bernstein, B. E. et al. An integrated encyclopedia of DNA elements in the human genome. Nature 489, 57–74 (2012).

57. Moriarity, B. S. et al. A Sleeping Beauty forward genetic screen identifies new genes and pathways driving osteosarcoma development and metastasis. Nat. Genet. 47, 615–24 (2015).

58. Lacruz, R. S. et al. Ontogeny of the maxilla in Neanderthals and their ancestors. Nat. Commun. 6, 8996 (2015).

59. Horvath, S. et al. The cerebellum ages slowly according to the epigenetic clock. Aging (Albany. NY). 7, 294–306 (2015).

60. Kulis, M. et al. Whole-genome fingerprint of the DNA methylome during human B cell differentiation. Nat. Genet. 47, 746–756 (2015).

61. Marco-Sola, S., Sammeth, M., Guigó, R. & Ribeca, P. The GEM mapper: fast, accurate and versatile alignment by filtration. Nat. Methods 9, 1185–1188 (2012).

62. Hansen, K. D., Langmead, B. & Irizarry, R. A. BSmooth: from whole genome bisulfite sequencing reads to differentially methylated regions. Genome Biol. (2012). doi:10.1186/gb-2012-13-10-R83

63. Dabney, J. et al. Complete mitochondrial genome sequence of a Middle Pleistocene cave bear reconstructed from ultrashort DNA fragments. Proc. Natl. Acad. Sci. U. S. A. 110, 15758–63 (2013).

64. Rohland, N. & Hofreiter, M. Ancient DNA extraction from bones and teeth. Nat. Protoc. 2, 1756– 1762 (2007).

65. Boyle, P. et al. Gel-free multiplexed reduced representation bisulfite sequencing for large-scale DNA methylation profiling. Genome Biol. 13, R92 (2012).

66. Barnett, R. & Larson, G. A phenol-chloroform protocol for extracting DNA from ancient samples. Methods Mol. Biol. 840, 13–19 (2012).

67. Fortin, J.-P., Triche, T. & Hansen, K. Preprocessing, normalization and integration of the Illumina HumanMethylationEPIC array. bioRxiv 065490 (2016). doi:10.1101/065490

68. Hernando-Herraez, I. et al. Dynamics of DNA Methylation in Recent Human and Great Ape Evolution. PLOS Genet 9, e1003763 (2013).

69. Emonet, E. G. & Kullmer, O. Variability in premolar and molar root number in a modern population of pan troglodytes verus. Anat. Rec. 297, 1927–1934 (2014).

70. Sirak, K. A. et al. A minimally-invasive method for sampling human petrous bones from the cranial base for ancient DNA analysis. Biotechniques 62, 283–289 (2017).

71. Haak, W. et al. Massive migration from the steppe was a source for Indo-European languages in Europe. Nature 522, 207–211 (2015).

72. Gokhman, D., Meshorer, E. & Carmel, L. Epigenetics: It’s Getting Old. Past Meets Future in Paleoepigenetics. Trends Ecol. Evol. 31, 290–300 (2016).

73. Hanghøj, K., et al. Fast, Accurate and Automatic Ancient Nucleosome and Methylation Maps with epiPALEOMIX. Mol. Biol. Evol. 33, 3284–3298 (2016).

74. Huang, D. W., Lempicki, R. a & Sherman, B. T. Systematic and integrative analysis of large gene lists using DAVID bioinformatics resources. Nat. Protoc. 4, 44–57 (2009).

75. Blasi, D. E. et al. Human sound systems are shaped by post-Neolithic changes in bite configuration. Science (80-.). (2019). doi:10.1126/science.aav3218

76. Guo, M. et al. Epigenetic profiling of growth plate chondrocytes sheds insight into regulatory genetic variation influencing height. Elife 6, (2017).

77. Bagheri-Fam, S. et al. Long-range upstream and downstream enhancers control distinct subsets of the complex spatiotemporal Sox9 expression pattern. Dev. Biol. 291, 382–397 (2006).

78. Sekido, R. & Lovell-Badge, R. Sex determination involves synergistic action of SRY and SF1 on a specific Sox9 enhancer. Nature 453, 930–934 (2008).

79. Varanasi, S. S. et al. Skeletal site-related variation in human trabecular bone transcriptome and signaling. PLoS One 5, e10692 (2010).

80. Martynoga, B. et al. Epigenomic enhancer annotation reveals a key role for NFIX in neural stem cell quiescence. Genes Dev. 27, 1769–1786 (2013).

81. Waki, H. et al. Global mapping of cell type-specific open chromatin by FAIRE-seq reveals the regulatory role of the NFI family in adipocyte differentiation. PLoS Genet. 7, e1002311 (2011).

