## Extended Data Table 9 for "Genes Affecting Vocal and Facial Anatomy Went Through Extensive Regulatory Divergence in Modern Humans"

| **Extraction** | **Mg used** | **ug/mL** | **Volume in uL** | **Total ug** |
| --- | --- | --- | --- | --- |
| Extraction 1 | 50 | 1.28 | 115 | 0.1472 |
| Extraction 2 | 100 | 2.51 | 115 | 0.2886 |
| Extraction 3 | 100 | 1.8 | 115 | 0.2162 |
| Blank control 1 | n/a | <0.010 ug/mL | 115 | 0 |
| Blank control 2 | n/a | <0.010 ug/mL | 115 | 0 |

**Extended Data Table 9. DNA extractions from chimpanzee femurs.**
