## Extended Data Table 11 for "Genes Affecting Vocal and Facial Anatomy Went Through Extensive Regulatory Divergence in Modern Humans"

| **Animal ID** | **Species** | **Sex** | **Age (yr)** | **Array ID** | **Position** |
| --- | --- | --- | --- | --- | --- |
| 4X0523 | chimpanzee | male | 10.20 | 200670680029 | R02C01 |
| 4X0406 | chimpanzee | male | 10.26 | 200665030025 | R07C01 |
| 4X0387 | chimpanzee | male | 13.46 | 200705360064 | R07C01 |
| 4-0191 | chimpanzee | unknown | unknown | 200705860055 | R05C01 |

**Extended Data Table 11.** Chimpanzee methylation array samples.
