## Extended Data Table 12 for "Genes Affecting Vocal and Facial Anatomy Went Through Extensive Regulatory Divergence in Modern Humans"

| **Region** | **Forward primer** | **Reverse primer** |
| --- | --- | --- |
| NFIX_1 | GGGATTGGAGAATTTGTTTTTGT | AAAAAATATCAACCCTTCCC |
| NFIX_2 | TTTTTGTTGATTAAAGGGGT | ATACCCTTAAACACAAAACT |
| NFIX_3 | TTTTTATTGGGAAGGGTTAA | CCTCCCAAAAAAAAAAAAAAC |
| NFIX_4 | TTTTTTGAGATTAGGTAGGGAA | TCACACCCACAATAAAAATAAAC |
| NFIX_5 (chimp) | GGAAGATAGGTTTTTTTTTTTAT | AAAAATCAACCTTTACCTTCATC |
| NFIX_5 (human) | GGAAGATAGGTTTTTTTTTTTGT | AAAAATCAACCTTTACCTTCATC |

**Extended Data Table 12. Primers used for Bisulfite-PCR**
